# Scale invariance during bacterial reductive division observed by an extensive microperfusion system

**DOI:** 10.1101/2020.06.25.171710

**Authors:** Takuro Shimaya, Reiko Okura, Yuichi Wakamoto, Kazumasa A. Takeuchi

## Abstract

In stable environments, cell size fluctuations are thought to be governed by simple physical principles, as suggested by recent finding of scaling properties. Here we show, using E. coli, that the scaling concept also rules cell size fluctuations under time-dependent conditions, even though the distribution changes with time. We develop a microfluidic device for observing dense and large bacterial populations, under uniform and switchable conditions. Triggering bacterial reductive division by switching to non-nutritious medium, we find that the cell size distribution changes in a specific manner that keeps its normalized form unchanged; in other words, scale invariance holds. This finding is underpinned by simulations of a model based on cell growth and intracellular replication. We also formulate the problem theoretically and propose a sufficient condition for the scale invariance. Our results emphasize the importance of intrinsic cellular replication processes in this problem, suggesting different distribution trends for bacteria and eukaryotes.

## Introduction

Recent studies on microbes in the steady growth phase suggested that the cellular body size fluctuations may be governed by simple physical principles. For instance, Giometto *et al*. (***Giometto et al. (2013)***) proposed that size fluctuations of various eukaryotic cells are governed by a common distribution function, if the cell sizes of a given species are normalized by their mean value (see also (***Zaoli et al. (2019)***)). In other words, the distribution of cell volumes *ν, p*(*v*), can be described as follows:

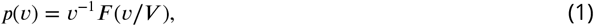

with a function *F*(·) and *V* = 〈*v*〉 being the mean cell volume. This property of distribution is often called scale invariance. Interestingly, this finding can account for power laws of community size distributions, i.e., the size distribution of all individuals regardless of species, which were observed in various natural ecosystems (***Camacho and Solé (2001)***; ***Marquet et al. (2005)***). Scale invariance akin to ***Equation 1*** was also found for bacteria (***Iyer-Biswas et al. (2014b)***; ***Kennard et al. (2016)***) for each cell age, and the function *F*(·) was shown to be robust against changes in growth conditions such as the temperature.

Those results, as well as theoretical models proposed in this context (***italic>Giometto et al. (2013)***; ***Iyer-Biswas et al. (2014a)***), have been obtained under steady environments, for which our understanding of single-cell growth statistics has also been significantly deepened recently (***Ho et al. (2018)***; ***Jun et al. (2018)***; ***Cadart et al. (2019)***). By contrast, it is unclear whether such a simple concept as scale invariance is valid under time-dependent conditions, where different regulations of cell cycle kinetics may come into play in response to environmental variations. In particular, when bacterial cells enter the stationary phase from the exponential growth phase, they undergo reductive division, during which both the typical cell size and the amount of DNA per cell decrease (***Nyström (2004)***; ***Kaprelyants and Kell (1993)***; ***Arias et al. (2012)***; ***Gray et al. (2019)***). Although this behavior itself is commonly observed in test tube cultivation, little is known about single-cell statistical properties during the transient. The bacterial reductive division is therefore an important model case for studying cell size statistics under time-dependent environments and testing the robustness of the scale invariance against environmental changes.

It has been, however, a challenge to observe large populations of bacteria under uniform yet time-dependent growth conditions. For steady conditions, the Mother Machine (***Wang et al. (2010)***), which allows for tracking of bacteria trapped in short narrow channels, was proved to be a powerful tool for measuring cell size statistics. In such experiments, the channel width needs to be adapted to cell widths in a given condition, and this renders the application to time-dependent conditions difficult. If we enlarge the channels, depletion of nutrients in deeper regions of the channels induces spatial heterogeneity, as discussed in ref.(***Cho et al. (2007)***; ***Mather et al. (2010)***) and later in this article. Hence, it is also required to develop a system that can uniformly control non-steady environments and give a sufficient amount of single-cell statistics with high efficiency.

In this study, we establish a novel microfluidic device, which we name the “extensive microperfusion system” (EMPS). This system can trap dense bacterial populations in a wide quasi-two-dimensional region and uniformly control the culture condition for a long time. We confirm that bacteria can freely swim and grow inside, and evaluate the uniformity and the switching efficiency of the culture condition. Then we use this system for quantitative observations of bacterial reductive division processes. We observe *Escherichia coli* cells and find that the distribution of cell volumes, collected irrespective of cell ages, maintain the scale invariance as in ***Equation 1*** at each time, with the mean cell size that gradually decreases. To obtain theoretical insights on this experimental finding, we devise a cell cycle model describing reductive division processes, by extending the single-cell Cooper-Helmstetter model (***Cooper and Helmstetter (1968)***; ***Wallden et al. (2016)***) for steady growth environments. We numerically find that this model indeed shows the scale invariance, confirming the robustness of this property. Further, we provide theoretical descriptions on the time evolution of the cell size distribution, and propose a condition for the scale invariance.

## Results

### Development of the extensive microperfusion system

In order to achieve uniformly controlled environments with dense bacterial suspensions, we adopt a perfusion system as developed in ref. (***Inoue et al. (2001)***). This system allows for supplying fresh medium through a porous membrane attached over the entire observation area, instead of supplying from an open end as polydimethylsiloxane (PDMS)-based systems usually do. In its prototypical setup, bacteria are confined in microwells made on a coverslip, covered by a cellulose porous membrane attached to the coverslip via biotin-streptavidin bonding. The pore size of the membrane is chosen so that it can confine bacteria and also that it can exchange nutrients and waste substances across the membrane. To continuously perfuse the system with fresh medium, a PDMS pad with a bubble trap is attached above the membrane by a two-sided frame seal (***Figure 1a*** and ***Figure 1–Figure Supplement 1a***). This setup can maintain a spatially homogeneous environment for cell populations in each microwell, in particular if the microwells are sufficiently shallow so that all cells remain near the membrane. However, because the soft cellulose membrane may droop and adhere to the bottom for wide and shallow microwells, the horizontal size of such quasi-two-dimensional microwells has been limited up to a few tens of micrometers, preventing from characterization of the instantaneous cell size distribution.

**Figure 1.**
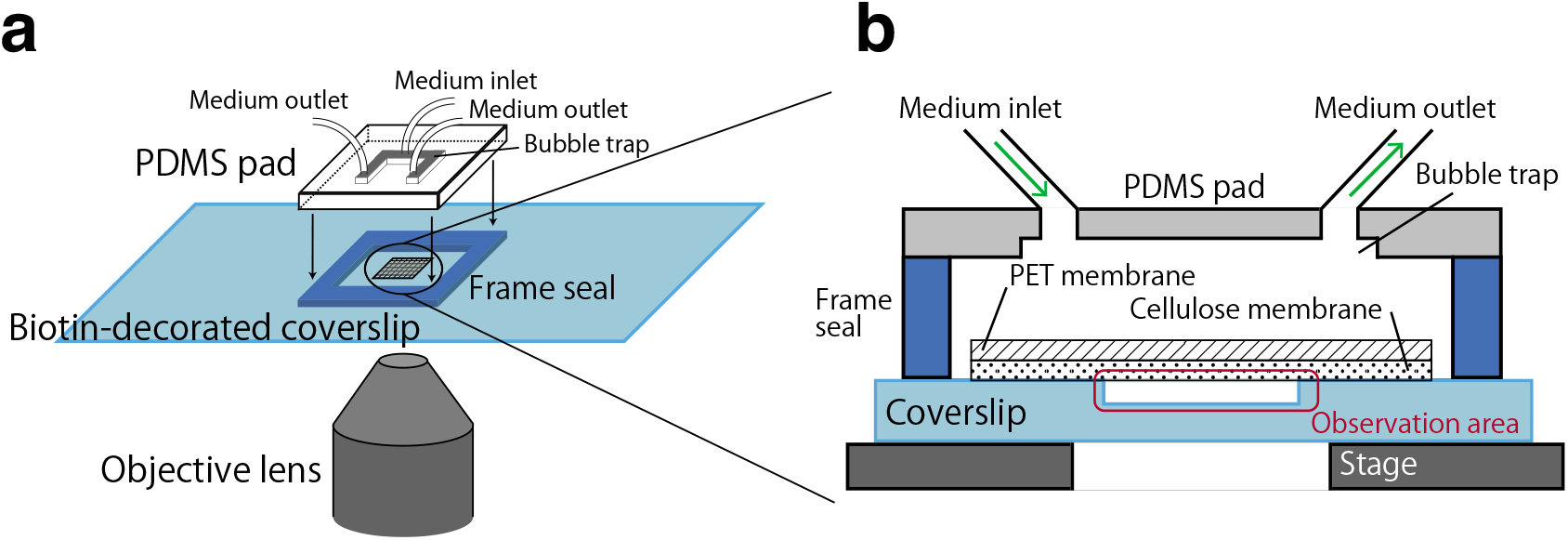
Sketch of the extensive microperfusion system (EMPS). **a** Entire view of the device. Microwells are created on a glass coverslip. We attach a PDMS pad on the coverslip with a square frame seal to fill the system with liquid medium. **b** Cross-sectional view inside the PDMS pad. A PET-cellulose bilayer porous membrane is attached via the biotin-streptavidin bonding. **Figure 1–Figure supplement 1.** Supplementary figures on the setup of the EMPS.

By the EMPS, we overcome this problem and realize quasi-two-dimensional wells sufficiently large for statistical characterization of cell populations. This is made possible by introducing a bilayer membrane, where the cellulose membrane is sustained by a polyethylene terephthalate (PET) porous membrane via biotin-streptavidin bonding (***Figure 1b, Figure 1–Figure Supplement 1b*** and Materials and Methods). Because the PET membrane is more rigid than the cellulose membrane, we can realize extended area without bending of the membrane. Specifically, in our setup with circular wells of 110 *μ*m diameter and 1.1 *μ*m depth, although a cellulose membrane alone is bent and adheres to the bottom of the well (***Figure 1–Figure Supplement 1c*** and ***Video 1***), our PET-cellulose bilayer membrane keeps flat enough so that bacteria can freely swim in the shallow well (***Figure 1–Figure Supplement 1d*** and ***Video 2***). The EMPS can realize such observations for a long time with little hydrodynamic perturbation by medium flow and no mechanical stress, which may exist in a PDMS-based device that holds cells mechanically. We also check whether the additional PET membrane may affect the growth condition of cells, by using *E. coli* MG1655 and M9 medium with glucose and amino acids (Glc+a.a.). We find that the doubling time of the cell population is 59 ± 10 min, which is comparable to that in the previous system without the PET membrane (***Inoue et al. (2001)***; ***Wakamoto et al. (2005)***; ***Hashimoto et al. (2016)***). Therefore, our bilayer membrane can still exchange medium efficiently.

**Video 1.** Growth of motile *E. coli* RP437 inside a well covered only by a cellulose membrane. The diameter of the well is 110 *μ*m, and the depth is 1.1 *μ*m. Being pressed by the bent cellulose membrane, cells do not swim but form clusters, extending toward the wall. The movie is played at 600× real-time speed.

**Video 2.** Growth of motile *E. coli* RP437 inside a well covered by a PET-cellulose bilayer membrane. The diameter of the well is 110 *μ*m, and the depth is 1.1 *μ*m. Cells freely swim inside the quasi-two-dimensional well. The movie is played at real-time speed.

**Video 3.** Coherent flow of non-motile bacterial cells driven by self-replication in a U-shape trap of the PDMS-based device. The trap is 30 *μ*m wide, 88 *μ*m long, and 1.1 *μ*m deep. *E.coli* strain W3110 ΔfliC Δflu ΔfimA is used.

**Video 4.** Coherent flow of non-motile bacterial cells driven by self-replication in a U-shape trap of the EMPS. The trap is 30 *μ*m wide, 80 *μ*m long, and 1.0 *μ*m deep. *E.coli* strain W3110 ΔfliC Δflu ΔfimA is used.

We then test the spatial uniformity of the culture condition. We design U-shape traps with an open end, for both the EMPS (***Figure 2a***) and for the conventional PDMS-based device (***Figure 2–Figure Supplement 1ab***). With this geometry, nutrients are supplied via diffusion from the open end in the PDMS-based system, while nutritious medium is directly and uniformly delivered through the membrane above the trap in the EMPS. When we culture *E. coli* MG1655 in M9(Glc+a.a.), the trap is eventually filled with cells, and they exhibit coherent flow toward the open end due to the cell growth and proliferation (***Figure 2b and Video 3,4***). To evaluate the uniformity of the cell growth, we measure the velocity field of the cell flow by particle image velocimetry (PIV) (***Figure 2–Figure Supplement 1cd***). The velocity component along the stream-wise direction (the *y* axis in ***Figure 2b***) averaged over the span-wise direction, *u*(*y*), clearly shows that the velocity profile is stable for the EMPS over long time periods, while it gradually decreases for the PDMS-based device (***Figure 2c*** Main Panel and Inset, respectively). The cell growth rate is then obtained by 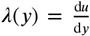, which is shown in ***Figure 2d***. The result shows that the growth rate *λ* is indeed uniform and kept constant for the EMPS (***Figure 2d***, Main Panel), while for the PDMS device it is heterogeneous, being higher near the open end (***Figure 2d***, Inset), and it decreases as time elapses. The growth rate decays at the distance of roughly 30 *μ*m from the open end, which is located near *y* ≈ 88 *μ*m for the PDMS device. This observation is consistent with the nutrient depletion length we evaluate by following the calculation by Mather et al. (***Mather et al. (2010)***), 32 *μ*m, for which we used the diffusion constant of glucose (***Marucci et al. (2006)***) and the division time of 60 min. Such heterogeneity is not seen in the EMPS. While cell growth regulation pathways may also be influenced by such factors as mechanical pressure caused by cell elongation (***Volfson et al. (2008)***; ***Boyer et al. (2011)***; ***Si et al. (2016)***; ***Chu et al. (2018)***), quorum sensing (***Bruger and Waters (2016)***; ***Ha et al. (2018)***), etc., our results indicate that the EMPS can indeed realize a uniform and stable culture condition while the same medium is kept supplied.

**Figure 2.**
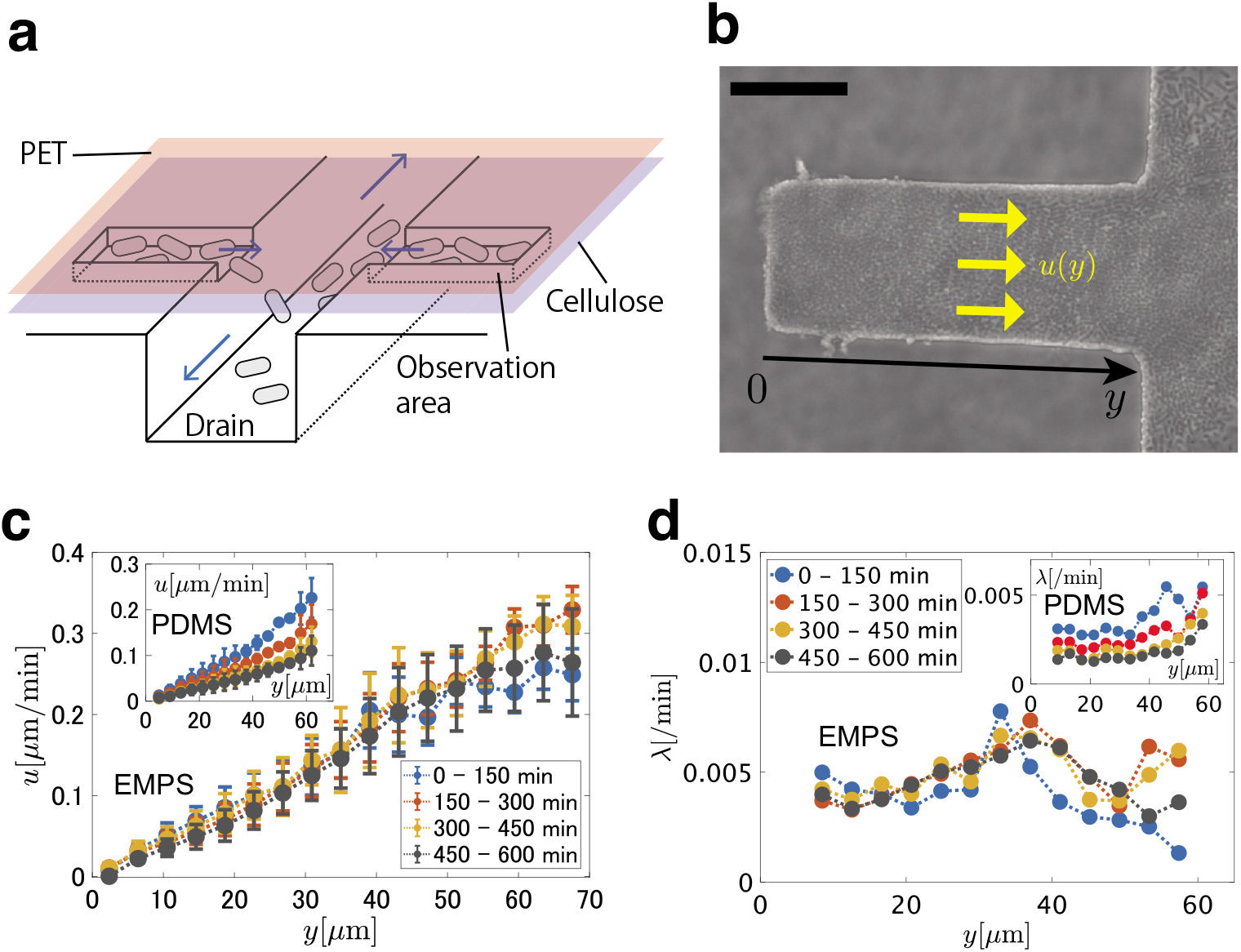
Cell growth measurements in U-shape traps in EMPS. **a** Sketch of the design of microchannels. Non-motile cells are trapped in the shallow observation area. Cells in the trap can escape to the deep drain channel. **b** Top view of the trap (30 *μ*m× 80 *μ*m, 1.0 *μ*m depth) filled with *E. coli* W3110 ΔfliC Δflu ΔfimA(see also ***Video 4***). The scale bar is 25 *μ*m. Coherent flow of cells driven by cell proliferation is directed toward the drain (15 *μ*m depth). **c** The stream-wise (*y*) component of the velocity field averaged over the span-wise direction (the two-dimensional velocity field is shown in ***Figure 2–Figure Supplement 1c***), *u*, measured in different time periods. The data were taken from a single trap. *t* = 0 is the time at which the trap is filled with cells. Error bars represent the standard deviation of the ensemble. The open end is located near *y* ≈ 81 *μ*m. (Inset) The same quantity measured for a PDMS-based device with a similar trap (***Figure 2–Figure Supplement 1abc***). The open end is at *y* ≈ 88 *μ*m. Time dependence is clearly observed. **d** The growth rate profiles *λ*, evaluated by 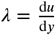, for the EMPS (main panel) and the PDMS-based device (inset). **Figure 2–Figure supplement 1.** Supplementary figures on the cell growth measurement. **Figure 2-source data 1.** Averaged velocity *u* and the standard deviation.

Another advantage of the EMPS is that we can also switch the culture condition, by changing the medium to supply. Here we evaluate how efficiently the medium in the well is exchanged. In the presence of non-motile *E. coli* W3110 ΔfliC Δflu ΔfimA, we switch the medium to supply from phosphate buffered saline (PBS) to PBS solution of rhodamine fluorescent dye, and monitor the fluorescent signal in a cross-section of the device by a confocal microscope (***Figure 3a***, from right to left). The result shows that the medium inside the well is exchanged uniformly in space (***Figure 3b***) and that it is almost completed within 2-4 min (***Figure 3d***). We also change the medium from PBS with rhodamine to that without rhodamine (***Figure 3a***, from left to right). The exchange then took longer time, *>* 5 min, presumably because of adsorption of rhodamine on the substrate and membrane (see ***Figure 3a***). In any case, the time to take for exchanging medium is much shorter than the timescale of the bacterial cell cycle. Our observations also indicate that the membrane is indeed kept flat above the well (***Figure 3a***) and that the Brownian motion of non-motile cells is hardly affected by relatively strong medium flow above the membrane (estimated at roughly 6 mm/sec) induced when switching the medium (***Video 5*** and ***Video 6***). We therefore conclude that the EMPS is indeed able to change the growth condition for cells under observation uniformly, without noticeable fluid flow perturbations.

**Figure 3.**
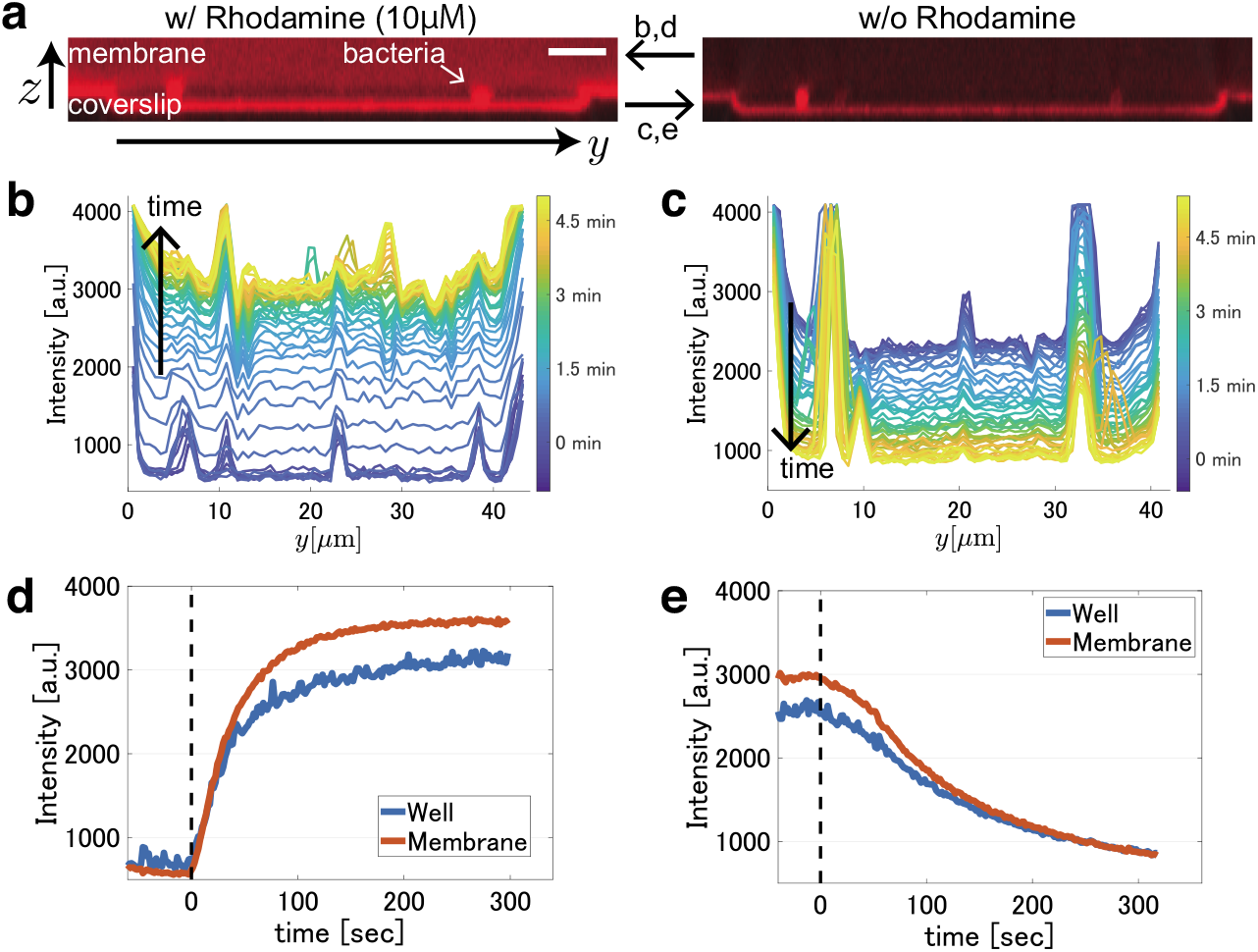
Direct observations of medium exchange in the EMPS. **a** Cross-sectional images of the device taken by confocal microscopy. The scale bar is 5 *μ*m. Medium flows from the front side to the rear side of the images. Although the flow speed of the medium is relatively high above the membrane, the diffusive motion of cells is hardly affected (see Movies S5 and S6). (left) A snapshot of the device filled with a PBS solution with 10 *μ*M rhodamine. Bacterial cells (W3110 ΔfliC Δflu ΔfimA) and the bilayer membrane are dyed and visualized. (right) A snapshot of the device filled with PBS without rhodamine. The surface of the coverslip and the cells still exhibit fluorescence because of adsorption of rhodamine. **b,c** Time evolution of the spatial profile of the fluorescent intensity, when the medium is switched to the rhodamine solution (**b**, see also ***Video 5***) and to the PBS without rhodamine (**c**, see also ***Video 6***). The intensity averaged over 5 pixels (0.6 *μ*m) from the substrate bottom is shown. Note that the location of the substrate bottom was detected by image analysis in each frame, in order to avoid the influence of vibrations (see Movies S5 and S6) caused by the high flow rate used here. The peaks seen in the profiles are due to bacterial cells, walls or dust. **d,e** Time series of the spatially averaged fluorescence intensity when the medium is switched to the rhodamine solution (**d**) and to the PBS without rhodamine (**e**). During the experiment, medium flowed above the membrane at a constant speed of approximately 6 mm/sec. *t* = 0 is the time at which the rhodamine solution entered the device (black dashed line). The spatial average of intensity in the well (blue curves) was taken in a square ROI of height 5 pixels (0.6 *μ*m) from the substrate bottom, and width 200 pixels (24 *μ*m) along the y-axis, around the center of the well. The spatial average of intensity in the membrane (red curves) was taken in a linear ROI of length 200 pixels (24 *μ*m) along the y-axis, located at 4.8 *μ*m above the substrate bottom, around the center. **Figure 3-source data 1.** Time series of averaged intensity.

**Video 5.** A cross-sectional movie of the EMPS, recorded while medium is switched from transparent PBS to a PBS solution of 10 *μ*M rhodamine. The diameter of the well is 45 *μ*m and the depth is 1.1 *μ*m. Afew non-motile *E.coli* (W3110 ΔfliC Δflu ΔfimA) are present in the well. The rhodamine solution flowed at a constant speed of approximately 6 mm/sec above the membrane (flow rate 60 ml/hr). The movie is played at 19× real-time speed.

**Video 6.** A cross-sectional movie of the EMPS, recorded while medium is switched from a PBS solution of 10 *μ*M rhodamine to transparent PBS. The diameter of the well is 45 *μ*m and the depth is 1.1 *μ*m. Afew non-motile *E.coli* (W3110 ΔfliC Δflu ΔfimA) are present in the well. The PBS without rhodamine flowed at a constant speed of approximately 6 mm/sec above the membrane (flow rate 60 ml/hr). The movie is played at 19× real-time speed.

### Characterization of bacterial reductive division by EMPS

Now we observe the reductive division of *E. coli* MG1655 in the EMPS, triggering starvation by switching from nutritious medium to non-nutritious buffer. In the beginning, a few cells are trapped in a quasi-two-dimensional well (diameter 55 *μ*m and depth 0.8 *μ*m) and grown in nutritious medium, until a microcolony composed of approximately 100 cells appear. We then quickly switch the medium to a non-nutritious buffer, which is continuously supplied until the end of the observation (see Materials and Methods for more details). By doing so, we intend to remove various substances secreted by cells, such as autoinducers for quorum sensing and waste products, to reduce their effects on cell growth (***Carbonell et al. (2002)***; ***Bruger and Waters (2016)***; ***Ha et al. (2018***); ***Maier and Pepper (2015)***). Throughout this experiment, the well is entirely recorded by phase contrast microscopy. We then measure the length and the width of all cells in the well, to obtain the volume *ν* of each cell by assuming the spherocylindrical shape, at different times before and after the medium switch. Here we mainly show the results for the case where the medium is switched from LB broth to PBS (denoted by LB → PBS) in ***Figure 4***, while the results for M9(Glc+a.a.) → PBS, and M9 medium with glucose (Glc) → M9 medium with *α*-methyl-D-glucoside (*α*MG), a glucose analog which cannot be metabolized (***Chou et al. (1994)***), are also presented in ***Figure 4–Figure Supplement 1*** and S5. We observe that, after switching to the non-nutritious buffer, the growth of the total volume decelerates (***Video 7, Video 8, Video 9, Figure 4b, Figure 4–Figure Supplement 1b*** and ***Figure 4–Figure Supplement 2b***), and the mean cell volume rapidly decays because of excessive cell divisions (***Video 7, Video 8, Video 9, Figure 4ac***). The volume change is mostly due to the decrease of the cell length, while we notice that the mean cell width may also change slightly (***Figure 4–Figure Supplement 3***). We consider that this is not due to osmotic shock (***Rojas et al. (2014)***), because then the cell width would increase when the osmotic pressure is decreased, which is contradictory to our observation for LB → PBS (***Table 2*** and ***Figure 4–Figure Supplement 3***). Such a change in cell widths was also reported for a transition between two different growth conditions (***Harris and Theriot (2016)***). In any case, ***Figure 4d*** shows how the distribution of the cell volumes *ν*, *p*(*ν,t*), changes over time; as the mean volume decreases, the histograms shift leftward and become sharper. However, when we take the ratio *ν/V*(*t*), with *V*(*t*) = (*ν*) being the mean cell volume at each time *t*, and plot *νp*(*ν, t*) instead, we find that all those histograms overlap onto a single curve (***Figure 4e***). In other words, we find that the time-dependent cell size distribution during the reductive division maintains the following scale-invariant form all the time:

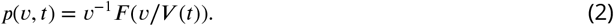

**Figure 4.**
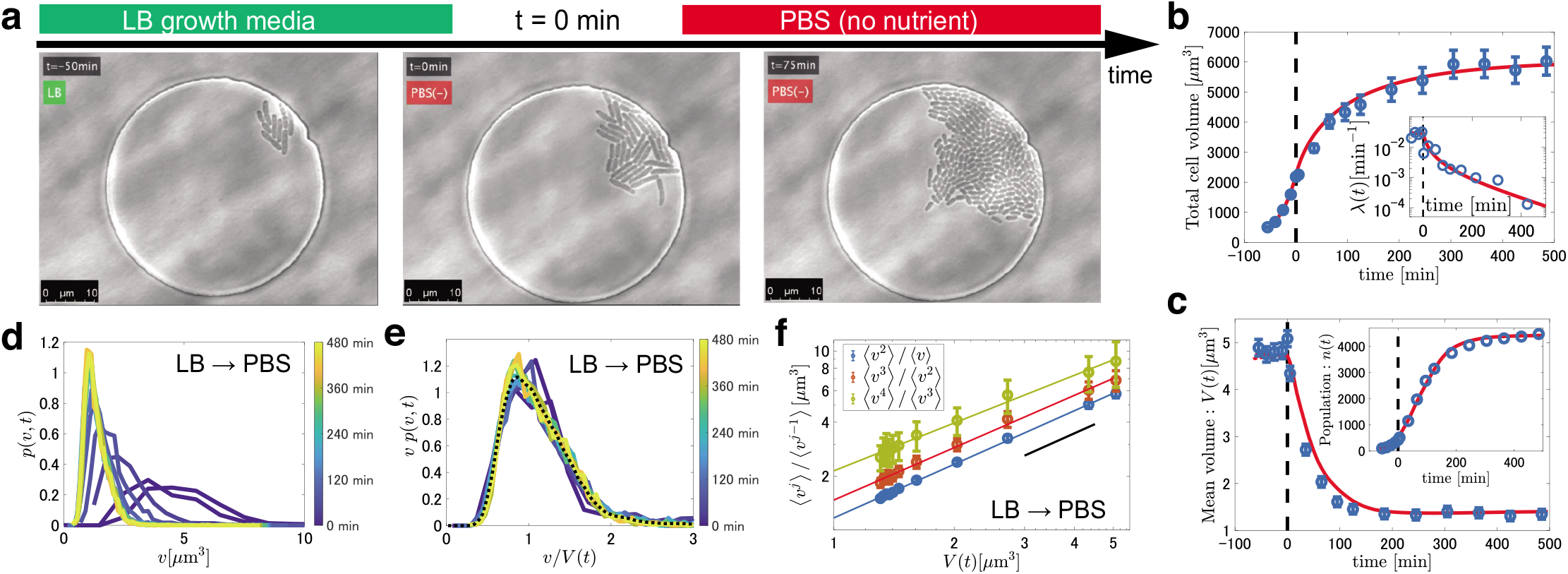
Results from the observations of reductive division. **a** Snapshots taken during the reductive division process of *E. coli* MG1655 in the EMPS. The medium is switched from LB broth to PBS at *t* = 0. See also ***Video 7*. b,c** Experimental data (blue symbols) for the total cell volume *V*_tot_(*t*) (**b**), the growth rate *λ*(*t*) (**b**, Inset), the mean cell volume *V*(*t*) (**c**) and the number of the cells *n*(*t*) (**c**, Inset) in the case of LB → PBS, compared with the simulation results (red curves). The error bars indicate segmentation uncertainty in the image analysis (see Materials and Methods). *t* = 0 is the time at which PBS enters the device (black dashed line). The data were collected from 15 wells. **d** Time evolution of the cell size distributions during starvation in the case of LB → PBS at *t* = 0,5,30,60,90,120,180,240,300,360,420,480 min from right to left. The sample size is *n*(*t*) for each distribution (see **c**, Inset). **e** Rescaling of the data in **d**. The overlapped curves indicate the function *F*(*ν/V*(*t*)) in ***Equation 2***. The dashed line represents the time average of the datasets. **f** The moment ratio 〈*v^l^*〉/〈*v*^*l*-1^) against *V*(*t*) = 〈*v*〉. The error bars were estimated by the bootstrap method with 1000 realizations. The colored lines represent the results of linear regression in the log-log plots (see ***Table 1*** for the slope of each line). The black solid lines are guides for eyes indicating unit slope, i.e., proportional relation. **Figure 4–Figure supplement 1.** Results from the observations of reductive division in the case of M9(Glc+a.a.) → PBS. **Figure 4–Figure supplement 2.** Results from the observations of reductive division in the case of M9(Glc) → M9(*α*MG). **Figure 4–Figure supplement 3.** Time series of the mean cell length and the mean cell width during the starvation process. **Figure 4–Figure supplement 4.** Cell-to-cell fluctuations of the septum position. **Figure 4–source data 1.** A time series of a data set of the individual cell volume in each case.

**Table 1.** Statistical data on the size distributions obtained by the experiments and the simulations. The exponents *a* and the coefficients of determination *R*^2^ are obtained by fitting (*v^j^*)/(*v*^*j*-1^) = *c*(*v*)^*α*^ in the corresponding log-log plots (***Figure 3f, Figure 4–Figure Supplement 1d, Figure 4–Figure Supplement 2d*** and ***Figure 6–Figure Supplement 1ad***). The standard deviation parameter *σ* of the log-normal distribution is defined in ***Equation 3*** in the main text.

| Media | Slope ($\beta$) | | | $R^2$ values | | | $\sigma$ |
| --- | --- | --- | --- | --- | --- | --- | --- |
| | $\langle v^2 \rangle / \langle v \rangle$ | $\langle v^3 \rangle / \langle v^2 \rangle$ | $\langle v^4 \rangle / \langle v^3 \rangle$ | $\langle v^2 \rangle / \langle v \rangle$ | $\langle v^3 \rangle / \langle v^2 \rangle$ | $\langle v^4 \rangle / \langle v^3 \rangle$ | |
| LB → PBS | 0.99(4) | 0.97(5) | 0.88(8) | 0.999 | 0.994 | 0.990 | 0.34(1) |
| M9(Glc+a.a.) → PBS | 0.97(3) | 0.96(4) | 0.95(5) | 1.000 | 0.998 | 0.993 | 0.29(2) |
| M9(Glc) → M9(αMG) | 1.00(4) | 1.02(4) | 1.10(5) | 1.000 | 0.999 | 0.996 | 0.28(1) |
| Simulation (LB → PBS) | 1.01(3) | 1.02(7) | 1.02(10) | 1.000 | 1.000 | 0.999 | 0.25(1) |
| Simulation (M9(Glc+a.a.) → PBS) | 0.98(2) | 0.96(3) | 0.95(4) | 1.000 | 1.000 | 0.999 | 0.22(1) |

**Table 2.** Culture conditions, ingredients and osmotic pressure.

| Medium<br>(Osmotic pressure [Osm/kg]) | Ingredients | Concentration |
| --- | --- | --- |
| LB broth (Millar)<br>(0.38) | Tryptone | 1 wt% |
|  | Sodium Chloride | 1 wt% |
|  | Yeast extract | 0.5 wt% |
| M9(Glc+a.a.) medium<br>(0.21) | Disodium Phosphate (Anhydrous) | 0.68 wt% |
|  | Monopotassium Phosphate | 0.3 wt% |
|  | Sodium Chloride | 0.05 wt% |
|  | Ammonium Chloride | 0.1 wt% |
|  | Magnesium Phosphate | 2 mM |
|  | Calcium Chloride | 0.1 mM |
|  | Glucose | 0.2 wt% |
|  | MEM Amino Acids solution (M5550, Sigma) | 1 wt% |
| M9(Glc) medium<br>(0.22) | Disodium Phosphate (Anhydrous) | 0.68 wt% |
|  | Monopotassium Phosphate | 0.3 wt% |
|  | Sodium Chloride | 0.05 wt% |
|  | Ammonium Chloride | 0.1 wt% |
|  | Magnesium Phosphate | 2 mM |
|  | Calcium Chloride | 0.1 mM |
|  | Glucose | 0.2 wt% |
| M9(αMG) medium<br>(0.21) | Disodium Phosphate (Anhydrous) | 0.68 wt% |
|  | Monopotassium Phosphate | 0.3 wt% |
|  | Sodium Chloride | 0.05 wt% |
|  | Ammonium Chloride | 0.1 wt% |
|  | Magnesium Phosphate | 2 mM |
|  | Calcium Chloride | 0.1 mM |
|  | Alpha-methyl-D-glucopyranoside. | 0.2 wt% |
| TB medium<br>(0.21) | Tryptone | 1 wt% |
|  | Sodium Chloride | 0.5 wt% |
| PBS(-)<br>(0.27) | Potassium Dihydrogenphosphate | 0.02 wt% |
|  | Disodium phosphate (Anhydrous) | 0.115 wt% |
|  | Potassium Chloride | 0.02 wt% |
|  | Sodium Chloride | 0.8 wt% |

This is analogous to ***Equation 1*** previously reported for the steady growth condition, but here importantly the mean volume *V*(*t*) changes over time significantly (***Figure 4c***). To further test the scale invariance of the distribution, we plot the moment ratios (*v^j^*)/(*v*^*j*-1^) with *j* = 2,3,4 and found them proportional to (*v*). This is indeed the relation expected if the distribution satisfies ***Equation 2*** (***Giometto et al. (2013)***). Remarkably, for all starving conditions that we test, we find that the scale invariance robustly appears (***Figure 4f, Figure 4–Figure Supplement 1d, Figure 4–Figure Supplement 2d***, and ***Figure 5***). Our results therefore indicate that the scale invariance as in ***Equation 2***, which has been observed for steady conditions (***Giometto et al. (2013)***; ***Zaoli et al. (2019)***), also holds in non-steady reductive division processes of *E. coli* rather robustly.

**Figure 5.**
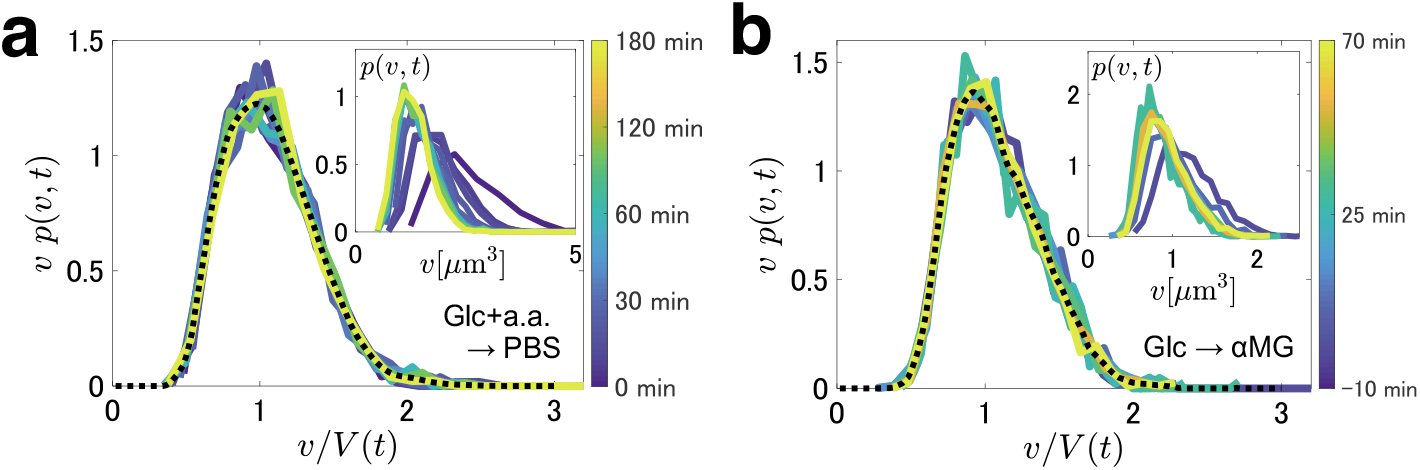
Rescaled cell size distributions. **a** The results for M9(Glc+a.a.) → PBS. The dashed line represents the time average of the datasets. The data were taken from 17 wells. The sample size ranges from *n*(0) = 685 to *n*(180) = 1260 (see ***Figure 4–Figure Supplement 1c***). (Inset) Time evolution of the non-rescaled cell size distributions at *t* = 0,10,20,30,40,50,60,90,120,180 min. **b** The results for M9(Glc) → M9(*α*MG). The dashed line represents the time average of the datasets. The data were taken from 26 wells. The sample size ranges from *n*(−5) = 1029 to *n*(65) = 1591 (see ***Figure 4–Figure Supplement 2c***). (Inset) Time evolution of the non-rescaled cell size distributions at *t* = −5,5,10,20,25,30,35,50,65 min. **Figure 5–Figure supplement 1.** Comparison of the function *F*(*v/V*(*t*)) = *vp*(*v,t*) among different cases.

**Video 7.** Reductive division of *E. coli* MG1655 for the case LB → PBS. The diameter of the well is 55 *μ*m and the depth is 0.8 *μ*m. Until *t* = 0, fresh LB broth was supplied at a constant flow speed of approximately 0.2 mm/sec above the membrane (flow rate 2 ml/hr). PBS entered the device at *t* = 0 and quickly replaced the LB broth, by setting a high flow speed ~ 6 mm/sec (60 ml/hr) until *t* = 5 min. After flushing, we continued supplying PBS at the flow speed of approximately 0.2 mm/sec (2 ml/hr).

**Video 8.** Reductive division of *E. coli* MG1655 for the case M9(Glc+a.a.) → PBS. The diameter of the well is 55 *μ*m and the depth is 0.8 *μ*m. PBS entered the device at *t* = 0. The flow rates were controlled and set in the same manner as for ***Video 7***.

**Video 9.** Reductive division of *E. coli* MG1655 for the case M9(Glc) → M9(*α*MG). The diameter of the well is 55 *μ*m and the depth is 0.8 *μ*m. M9(*α*MG) entered the device at *t* = 0. The flow rates were controlled and set in the same manner as for ***Video 7***.

In addition to the robustness of the scaling relation (***Equation 2***), the functional form of the scale-invariant distribution, i.e., that of *F*(*x*), is of central interest. In fact, we find that our observations for *E. coli* are significantly different from those for unicellular eukaryotes reported by Giometto *et al*. (***Giometto et al. (2013)***) (***Figure 5–Figure Supplement 1a***). More precisely, they showed that the rescaled cell size distribution for unicellular eukaryotes is well fitted by the log-normal distribution, which corresponds to

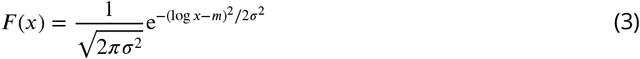

with *σ* = 0.471(3), and *m* = −*σ*^2^/2 due to the normalization 〈*x*〉 = 1. We find that our data for *E. coli* can also be fitted by a single log-normal distribution (***Figure 5–Figure Supplement 1a***, yellow dotted line), but here the value of *σ*, evaluated by the standard deviation of log *x*, is found to be *σ* = 0.304(6), much lower than *σ* = 0.471(3) for the unicellular eukaryotes. In the literature, a previous study on *B. subtilis* (***Wakita et al. (2010)***) reported values of *σ* from 0.24 to 0.26, which are comparable to our results shown in ***Table 1***. Compared to this substantial difference between bacteria and unicellular eukaryotes, the dependence on the environmental factors seems to be much weaker, as suggested by our observations under three different growth conditions (***Figure 5–Figure Supplement 1a***). Note, however, that we also noticed indications of weak dependence on the growth condition (***Figure 5–Figure Supplement 1b***). Clarifying the scope of the universality, or specifically, what factors affect the scale-invariant distribution and how strongly they do, is therefore an important open problem left for future studies.

### Modeling the reductive division process

To obtain theoretical insights on the experimentally observed scale invariance of the cell size distributions, we construct a simple cell cycle model for the bacterial reductive division. For the steady growth conditions, a large number of studies on *E. coli* have been carried out to clarify what aspect of cells triggers the division event ***Jun et al. (2018)***; ***Ho et al. (2018)***). Significant advances have been made recently to provide molecular-level understanding (***Ho et al. (2018)***; ***Ho and Amir (2015)***; ***Harris and Theriot (2016)***; ***Wallden et al. (2016)***; ***Micali et al. (2018a,b)***). Here we extend such a model to describe the starvation process.

One of the most established models in this context is the Cooper-Helmstetter (CH) model (***Cooper and Helmstetter (1968)***; ***Wang and Levin (2009)***), which consists of cellular volume growth and multifork DNA replication (***Figure 6a***). In this model, completion of the DNA replication triggers the cell division, and this gives a homeostatic balance between the DNA amount and the cell volume. An unknown factor of the CH model is how DNA replication is initiated, and a few studies attempted to fill this gap to complement the CH model (***Ho and Amir (2015)***; ***Wallden et al. (2016)***). Ho and Amir (***Ho and Amir (2015)***) assumed that replication is initiated when a critical amount of “initiators” accumulate at the origin of replication. In the presence of a constant concentration of autorepressors, expressed together with the initiators, this assumption means that the cellular volume increases by a fixed amount between two initiation events. In contrast, Wallden *et al*. (***Wallden et al. (2016)***) proposed another model named the single-cell CH (sCH) model, in which they considered, based on experimental observations, that the initiation occurs when the cell volume exceeds a given threshold independent of the growth rate and the initial volume of the cell. Albeit not exactly (***Micali et al. (2018a)***), both models successfully reproduced major characteristics of cell cycles in the exponential growth phase, such as the “adder” principle (***Campos et al. (2014)***; ***Taheri-Araghi et al. (2015)***; ***Jun et al. (2018)***). For modeling the reductive division process, we choose to extend the sCH model by Wallden *et al*., which requires a smaller number of assumptions to add to describe time-dependent processes studied here.

**Figure 6.**
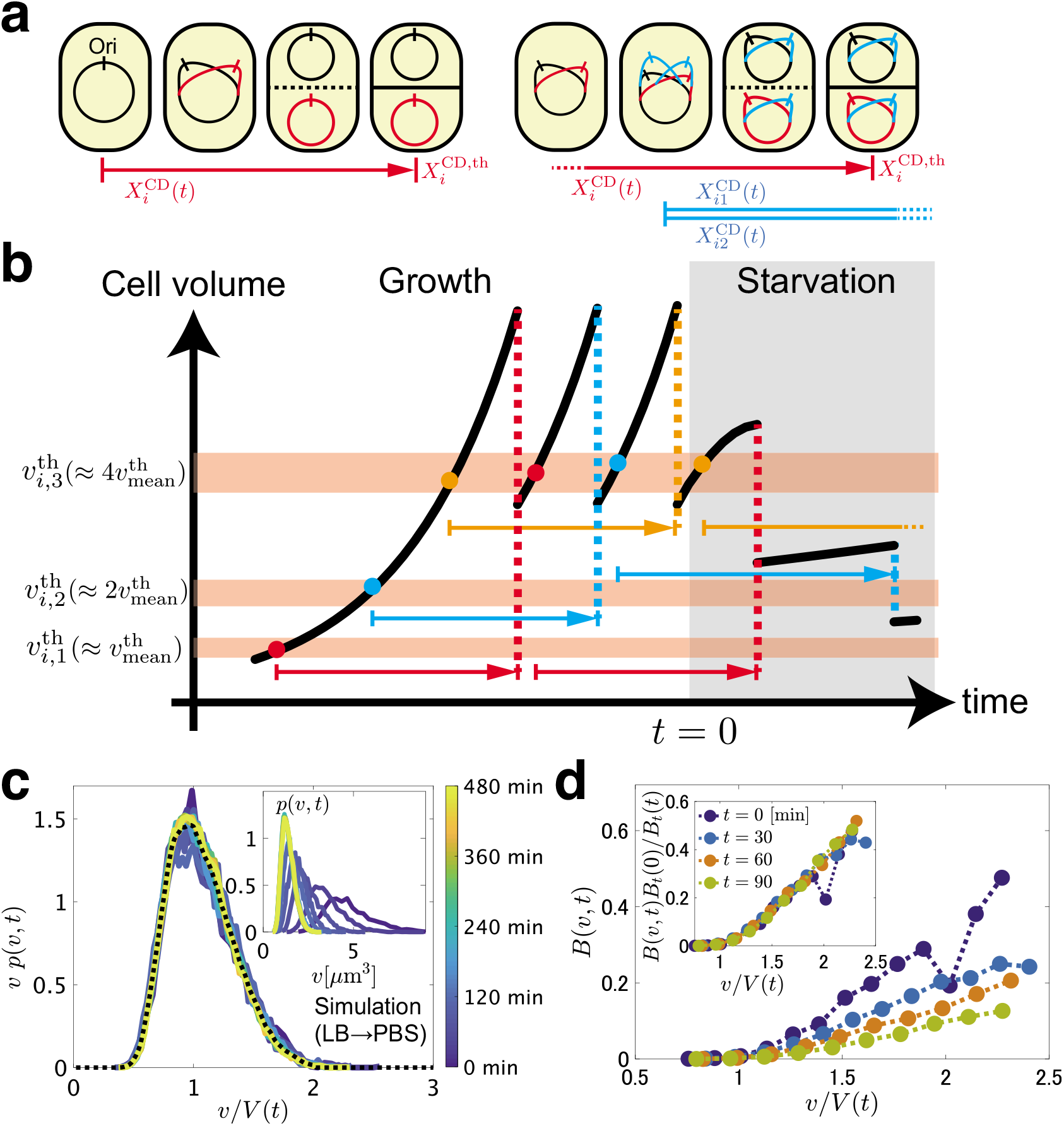
Model of reductive division and simulation results. **a** Single and multifork intracellular replication processes. Progress of each replication is represented by a coordinate 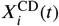, which ends at 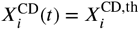 by triggering cell division. In the multifork process, all replications proceed simultaneously at the same rate *μ*(*t*). **b** Illustration of cell cycles in this model. Each colored arrow represents a single intracellular replication process, which starts when the cell volume *v_i_*(*t*) exceeds an initiation volume threshold 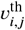. **c** Overlapping of the rescaled cell size distributions during starvation in the model for LB → PBS. The dashed line represents the time average of the datasets. (Inset) The non-rescaled cell size distributions at *t* = 0,5,30,60,90,120,180,240,300,360,420,480 min from right to left. **d** Numerically measured division rate, *B*(*v,t*), in the model for LB → PBS. See ***Appendix 2*** for the detailed measurement method. (Inset) Test of the condition of ***Equation 8***. Here *B_t_*(0)/*B_t_*(*t*) is evaluated by 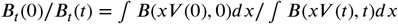, with *x* running in the range 0 ≤ *x* ≤ 1.8. Overlapping of the data demonstrates that ***Equation 8*** indeed holds in our model. **Figure 6–Figure supplement 1.** Supplementary figures on the simulation results. **Figure 6–Figure supplement 2.** Test of the self-consistent equations derived in ***Appendix 2***. **Figure 6–source data 1.** A time series of a data set of the individual cell volume obtained by the simulation (LB→PBS). **Figure 6–source data 2.** A time series of a data set of the individual cell volume obtained by the simulation (M9(Glc.+a.a.)→PBS).

The model consists of two processes that proceed simultaneously, namely the volume growth and the intracellular replication. The volume of each cell (indexed by *i*), *v_i_*(*t*), grows as 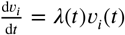, with a time-dependent growth rate *λ*(*t*). Following Wallden *et al*.’s sCH model (***Wallden et al. (2016)***), we assume that, when the cell volume *v_i_*(*t*) exceeds a threshold 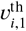 given below, DNA replication starts. This initiates the C+D period in the bacterial cell cycle (***Wang and Levin (2009)***; ***Cooper and Helmstetter (1968)***; ***Wallden et al. (2016)***), in which all processes regarding the cell division take place. Its progression is represented by a coordinate 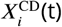, which starts form zero and increases at time-dependent speed 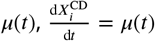. When 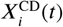 reaches a threshold 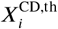, the cell divides (***Figure 6a***, left), leaving two daughter cells of volumes 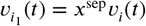 and 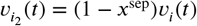. Here, *x*^sep^ is randomly drawn from the Gaussian distribution with mean 0.5 (see below for its standard deviation). It is also possible that the volume *v_i_*(*t*) reaches the second threshold 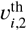, before this cell divides. Then the replications for the future daughter cells, 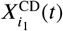 and 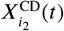, start and run simultaneously (***Figure 6a***, right and b). This, called the multifork replication (***Wang and Levin (2009)***; ***Cooper and Helmstetter (1968)***), is well-known for fast growing bacteria such as *E. coli* and *B. subtilis*. Similarly, at 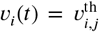, the replication for the *j*th generation is triggered. Following Wallden *et al*. (***Wallden et al. (2016)***), we assume that the initiation volume 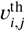 takes a constant value per chromosome, so that typically 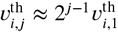. To take into account stochastic nature of division events, 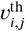 is generated randomly from the Gaussian distribution with mean 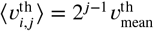 and standard deviation 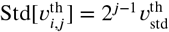. Similarly, 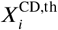 is also a Gaussian random variable with 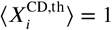 and 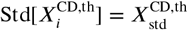. Here, 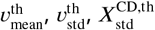 as well as 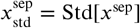 are considered to be parameters.

Now we are left to determine the two time-dependent rates, *λ*(*t*) and *μ*(*t*). Here we consider the situation where growth medium is switched to non-nutritious buffer at *t* = 0; therefore, *t* denotes time passed since the switch to the non-nutritious condition. First, we set the volume growth rate *λ*(*t*) on the basis of the Monod equation (***Monod (1949)***), assuming that substrates in each cell are simply diluted by volume growth and consumed at a constant rate, without uptake because of the non-nutritious condition considered here. As a result, we obtain

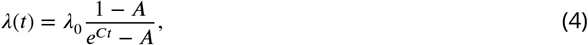

with constant parameters *A* and *C*, and the growth rate *λ*_0_(= *λ*(0)) in the exponential growth phase (see ***Appendix 1*** for details). For the replication speed *μ*(*t*), or more precisely the progression speed of the C+D period, we first note that the C+D period mainly consists of DNA replication, followed by its segregation and the septum formation (***Wang and Levin (2009)***). Most parts of those processes involve biochemical reactions of substrates, such as deoxynucleotide triphosphates for the DNA synthesis, and assembly of macromolecules such as FtsZ proteins for the septum formation. We therefore consider that the replication speed is determined by the intracellular concentration of relevant substrates and macromolecules, which is known to decrease in the stationary phase (***Buckstein et al. (2008)***; ***Sekar et al. (2018)***). By assuming an exponential decay of the substrate concentration, together with the Hill equation for the binding probability of ligands to receptors, we obtain the following equation for the replication speed:

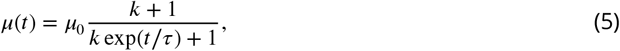

with parameters *k* and *τ*, and the replication speed *μ*_0_(= *μ*(0)) in the growth phase (see ***Appendix 1***).

The parameter values are determined from the experimentally measured total cell volume and the cell number, which our simulations turn out to reproduce very well (***Figure 4bc and Figure 4–Figure Supplement 1bc***), with the aid of relations reported by Wallden *et al*. (***Wallden et al. (2016)***) for some of the parameters (see ***Table 3*** for the parameter values used in the simulations, and Materials and Methods for the estimation method). With the parameters fixed thereby, we measure the cell size fluctuations at different times and find the scale invariance similar to that revealed experimentally (***Figure 6c*** and ***Figure 6–Figure Supplement 1bc***). The proportionality of the moment ratios is also confirmed (***Figure 6–Figure Supplement 1ad*** and ***Table 1***). Interestingly, the numerically obtained distribution is found to roughly reproduce the experimental one (***Figure 5–Figure Supplement 1a***), even though no information on the distribution is used for the parameter adjustment. Note that the scale invariance emerges despite the existence of characteristic scales in the model definition, such as the typical initiation volume 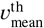. This suggests the existence of a statistical principle underlying the scale invariance, which is not influenced by details of the model and the experimental conditions.

**Table 3.** Parameters used for the simulations. The method of parameter determination is described in Materials and Methods in the main article.

|  | Parameters | LB→PBS | M9(Glc+a.a.)→PBS |
| --- | --- | --- | --- |
| Parameters on the exponential growth phase | $\lambda(0) = \lambda_0$ | $0.029 \text{ min}^{-1}$ | $0.010 \text{ min}^{-1}$ |
| | $\mu(0)^{-1} = \mu_0^{-1}$ | $1.3\lambda_0^{-0.84} + 42 \simeq 67 \text{ min}$ | $1.3\lambda_0^{-0.84} + 42 \simeq 104 \text{ min}$ |
| | $v_{\text{mean}}^{\text{th}}$ | $1.0 \mu\text{m}^3$ | $0.9 \mu\text{m}^3$ |
| | $v_{\text{std}}^{\text{th}}$ | $0.1 \times v_{\text{mean}}^{\text{th}} = 0.1 \mu\text{m}^3$ | $0.1 \times v_{\text{mean}}^{\text{th}} = 0.09 \mu\text{m}^3$ |
| | $X_{\text{std}}^{\text{CD,th}}$ | $0.05 \times \langle X_i^{\text{CD,th}} \rangle = 0.05$ | $0.05 \times \langle X_i^{\text{CD,th}} \rangle = 0.05$ |
| | $x_{\text{std}}^{\text{sep}}$ | $0.0325$ | $0.0325$ |
| Time-dependent rates | $\lambda(t) = \lambda_0 \frac{1-A}{e^{Ct} - A}$ | $A = 0.93, C = 0.0059 \text{ min}^{-1}$ | $A = 0.84, C = 0.011 \text{ min}^{-1}$ |
| | $\mu(t) = \mu_0 \frac{k+1}{ke^{t/\tau} + 1}$ | $k = 10, \tau = 75 \text{ min}$ | $k = 0.1, \tau = 24 \text{ min}$ |

### Theoretical conditions for the scale invariance

To seek for a possible mechanism leading to the scale invariance, here we describe, theoretically, the time dependence of the cell size distribution in a time-dependent process. Suppose *N*(*v,t*)*dv* is the number of the cells whose volume is larger than *v* and smaller than *v* + *dv*. If we assume, for simplicity, that a cell of volume *v* can divide to two cells of volume *v*/2, at probability *B*(*v,t*), we obtain the following time evolution equation:

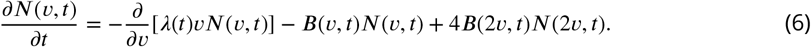

Note that this equation has been studied by numerous past studies for understanding stable distributions in steady conditions (***Sinko and Streifer (1971)***; ***Diekmann et al. (1983)***; ***Tyson and Diekmann (1986)***; ***Robert et al. (2014)***; ***Giometto et al. (2013)***; ***Hosoda et al. (2011)***), but here we explicitly include the time dependence of the division rate, *B*(*v,t*), for describing the transient dynamics. To clarify a condition for this equation to have a scale-invariant solution, here we assume the scale invariant form, ***Equation 2***, where *p*(*v,t*) = *N*(*v,t*)/*n*(*t*) and *n*(*t*) is the total number of the cells, and obtain the following self-consistent equation (see ***Appendix 2*** for derivation):

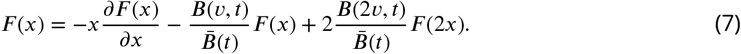

Here, *x* = *v*/*V*(*t*) and 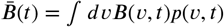. For the scale invariance, ***Equation 7*** should hold at any time *t*. This is fulfilled if *B*(*v,t*) can be expressed in the following form (see ***Appendix 2***):

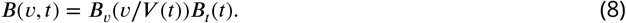

This is a sufficient condition for the cell size distribution to maintain the scale invariant form, ***Equation 2***, during the reductive division. It is important to remark that, as opposed to ***Equation 6, Equation 7*** does not include the growth rate *λ*(*t*) explicitly. The scale-invariant distribution *F*(*x*) is therefore completely characterized by the division rate *B*(*v, t*) in this framework.

To test whether the condition of ***Equation 8*** is satisfied in our model, we measure the division rate *B*(*v,t*) in our simulations (***Figure 6d***). The data overlap if *B*(*v,t*)*B_t_*(0)/*B_t_*(*t*) is plotted against *v*/*V*(*t*), demonstrating that ***Equation 8*** indeed holds here. In our model, the replication speed *μ*(*t*) is assumed to be given completely by the concentration of replication-related substrates, which takes the same value for all cells. This may be why the separation of variable, ***Equation 8***, effectively holds, and the scale invariance follows. On the other hand, our theory does not seem to account for the functional form *F*(*x*) of the scale-invariant distribution; the right hand side of ***Equation 7*** differs significantly from the left hand side, if the numerically obtained *B*(*v,t*) is used together with the function *F*(*x*) from the simulations or the experiments (***Figure 6–Figure Supplement 2***). The disagreement did not improve by taking into account the effect of septum fluctuations. The lack of quantitative precision is probably not surprising given the simplicity of the theoretical description, which incorporates all effects of intracellular replication processes into the simple division rate function *B*(*v,t*). The virtue of this theory is that it clarifies it is the replication process, not the cell body growth rate, that seems to have direct relevance in the scale invariance and the functional form of the cell size distribution. The significant difference in *F* (*x*) identified between bacteria and unicellular eukaryotes (***Figure 5–Figure Supplement 1a***) may be originated from the different replication mechanisms that the two taxonomic domains adopt.

### Concluding remarks

In this work, we developed a novel membrane-based microfluidic device that we named the extensive microperfusion system (EMPS), which can realize a uniformly controlled environment for wide-area observations of microbes. We believe that the EMPS has potential applications in a wide range of problems with dense cellular populations, including living active matter systems (***Bär et al. (2019)***; ***Be’er and Ariel (2019)***) and biofilm growth (***Hall-Stoodley et al. (2004)***; ***Boudarel et al. (2018)***; ***Fuqua et al. (2019)***). Here we focused on statistical characterizations of single cell morphology during the reductive division of *E. coli*. Thanks to the EMPS, we recorded the time-dependent distribution of cell size fluctuations and revealed that the rescaled distribution is scale-invariant and robust against the environmental change, despite the decrease of the mean cell size. This finding was successfully reproduced by simulations of a model based on the sCH model (***Wallden et al. (2016)***), which we propose as an extension for dealing with time-dependent environments. We further inspected theoretical mechanism behind this scale invariance and found the significance of the division rate function *B*(*v,t*). We obtained a sufficient condition for the scale invariance, ***Equation 8***, which was indeed confirmed in our numerical data.

After all, our theory suggests that mechanism of intracellular replication processes may have direct impact on the scale-invariant distribution, which may account for the significant difference we identified between bacteria and eukaryotes (***Figure 5–Figure Supplement 1a***). Since the number of species studied in each taxonomic domain is rather limited (*E. coli* (this work) and *B. subtilis* (***Wakita et al. (2010)***) for bacteria, 13 protist species for eukaryotes (***Giometto et al. (2013)***)), it is of crucial importance to test the distribution trend further in each taxonomic domain, and to clarify how and to what extent the cell size distribution is determined by the intracellular replication dynamics. We also note that the scale-invariant distribution *F*(*x*) might depend weakly on the culture condition in the exponential growth phase (***Figure 5–Figure Supplement 1b***). If this dependence on the growth condition is significant, we expect that conditions for the scale invariance are not met when switching between different growth environments. Investigations of cell size fluctuations in such cases, both experimentally and theoretically, will be an important step toward clarifying the scope of the universality of the scale-invariant cell size distributions. Combining with other theoretical methods, such as models considering the cellular age (***Grilli et al. (2017)***) and renormalization group approaches for living cell tissues (***Rulands et al. (2018)***), may be useful in this context. The influence of cell-to-cell interactions, e.g., quorum sensing (***Bruger and Waters (2016)***; ***Ha et al. (2018)***), may also be important. We hope that our understanding of the population-level response against nutrient starvation will be further refined by future experimental and theoretical investigations.

## Materials and Methods

### Strains and culture media

We used wild-type *E. coli* strains (MG1655 and RP437) and a mutant strain (W3110 ΔfliC Δflu ΔfimA) in this study. Culture media and buffer are listed in ***Table 2***. The osmotic pressure of each medium was measured by the freezing-point depression method by the OSMOMAT 030 (Genotec, Berlin Germany). Details on the strains and culture conditions in each experiment are provided below.

### Fabrication of the PDMS-based device

We prepared PDMS-based microfluidic devices by following the method reported in *Ref*.(***Nakaoka and Wakamoto (2017)***). We adopted a microchannel geometry similar to that in *Ref*.(***Mather et al. (2010)***), which consists of deep drain channels and shallow U-shape traps (***Figure 2–Figure Supplement 1ab***). In our setup, the drain channels are 25 *μ*m deep, and the U-shape traps are 30 *μ*m width, 70-90 *μ*m long, and 1.1 *μ*m deep. Tygon tubes of 1/16” outer diameter were connected to the inlets and the outlets (***Figure 2–Figure Supplement 1b***) via steel tubes (Elveflow, Paris France). The width of the drains (see ***Figure 2–Figure Supplement 1b***) and the length of tygon tubes were adjusted to realize the desired flow rate in the drain near the U-shape traps. The tube length was 55 cm for the medium inlet, 40 cm for the medium outlet, and 30 cm for the waste outlet. The cell inlet was connected to a syringe filled with bacterial suspension via a 10 cm tube, to inject cells at the beginning of observation. After the intrusion of cells, the syringe with the suspension was fixed, and no flow was generated around the cell inlet.

### Fabrication of the EMPS

The EMPS consists of a microfabricated glass coverslip, a bilayer porous membrane and a PDMS pad. The microfabricated coverslip and the PDMS pad were prepared according to ref. (***Inoue et al. (2001)***; ***Hashimoto et al. (2016)***). We fabricated the bilayer porous membrane by combining a streptavidin decorated cellulose membrane and a biotin decorated polyethylene-terephthalate (PET) membrane. The streptavidin decoration of the cellulose membrane (Spectra/Por 7, Repligen, Waltham Massachusetts, molecular weight cut-off 25000) was realized by the method described in ref. (***Inoue et al. (2001)***; ***Hashimoto et al. (2016)***). The PET membrane (Transwell 3450, Corning, Corning New York, nominal pore size 0.4 *μ*m) was decorated with biotin as follows. We soaked a PET membrane in 1 wt% solution of 3-(2-aminoethyl aminopropyl) trimethoxysilane (Shinetsu Kagaku Kogyo, Tokyo Japan) for 45 min, dried it at 125°C for 25 min and washed it by ultrasonic cleaning in Milli-Q water for 5 min. This preprocessed PET membrane was stored in a desiccator at room temperature, until it was used to assemble the EMPS.

The EMPS was assembled as follows. The preprocessed PET membrane was cut into 5 mm×5 mm squares, soaked in the biotin solution for 4 hours and dried on filter paper. The biotin decorated PET membrane was attached with a streptavidin decorated cellulose membrane, cut to the size of the PET membrane, by sandwiching them between agar pads (M9 medium with 2 wt% agarose). In the meantime, a 1 *μ*l droplet of bacterial suspension was inoculated on a biotin decorated coverslip (see also details below). We then took the bilayer membrane from the agar pad, air-dried for tens of seconds, and carefully put on the coverslip on top of the bacterial suspension. The bilayer membrane was then attached to the coverslip via streptavidin-biotin binding as shown in ***Figure 1–Figure Supplement 1b***. We then air-dried the membrane for a minute and attached a PDMS pad on the coverslip by a double-sided tape.

### Observation of motile *E. coli* in the EMPS

We used a wild-type motile strain of *E. coli*, RP437. First, we inoculated the strain from a glycerol stock into 2 ml TB medium (see ***Table 2*** for components) in a test tube. After shaking it overnight at 37 °C, we transferred 20 *μ*l of the incubated suspension to 2 ml fresh TB medium and cultured it until the optical density (OD) at 600 nm wavelength reached 0.1-0.5. The bacterial suspension was finally diluted to OD = 0.1 before it was inoculated on the coverslip of the EMPS.

Regarding the device, here we compared the EMPS with the previous system developed in ref. (***Inoue et al. (2001)***), whose membrane was composed of a cellulose membrane alone instead of the bilayer one. In either case, we used a substrate with circular wells of 110-210 *μ*m diameter and 1.1 *μ*m depth. After the assembly of the device with the bacterial suspension, it was fixed on the microscope stage inside an incubation box maintained at 37 °C. The microscope we used was Leica DMi8, equipped with a 63x (N.A. 1.30) oil immersion objective and operated by Leica LasX. To fill the device with medium, we injected fresh TB medium stored at 37 °C from the inlet (***Figure 1–Figure Supplement 1***), at the rate of 60 ml/hr for 5 min by a syringe pump (NE-1000, New Era Pump Systems, Farmingdale New York).

During the observation, TB medium was constantly supplied from the inlet at the rate of 2 ml/hr (flow speed above the membrane was approximately 0.2 mm/sec) by the syringe pump. Cells were observed by phase contrast microscopy and recorded at the time interval of 30 sec for the cellulose-membrane device (***Figure 1–Figure Supplement 1c***) and 118 msec for the EMPS (***Figure 1–Figure Supplement 1d***). The time interval for the former was long, because cells hardly moved in this case (see ***Video 1***). We checked that the EMPS can realize quasi-two-dimensional space in which bacteria can freely swim, even for large wells, at least up to 210 *μ*m in diameter, while we have not investigated the reachable largest diameter.

### Cell growth measurement in U-shape traps in the PDMS-based device

We used a non-motile mutant strain W3110 without flagella and pili (ΔfliC Δflu ΔfimA) to prevent cell adhesion to the surface of a coverslip. Before the time-lapse observation, we inoculated the strain from a glycerol stock into 2 ml M9 medium with glucose and amino acids (Glc+a.a.) (see also ***Table 2***) in a test tube. After shaking it overnight at 37 °C, we transferred 20 *μ*l of the incubated suspension to 2 ml fresh M9(Glc+a.a.) medium and cultured it until the OD at 600 nm wavelength reached 0.4-0.5. We then injected the bacterial suspension into the device from the cell inlet (***Figure 2–Figure Supplement 1b***) and left it until a few cells entered the U-shape traps. The device was placed on the microscope stage, in the incubation box maintained at 37 °C. The microscope we used was Leica DMi8, equipped with a 63x (N.A. 1.30) oil immersion objective and operated by Leica LasX.

During the observation, we constantly supplied M9(Glc+a.a.) medium and 0.5 wt% bovine serum albumin (BSA) from the medium inlet (***Figure 2–Figure Supplement 1b***) at the rate of 0.7 ml/hr (flow speed in the drain was approximately 1 mm/sec). Cells were observed by phase contrast microscopy and recorded at the time interval of 3 min. The velocity field of the coherent flow was obtained by particle image velocimetry, using MatPIV (MATLAB toolbox). The stream-wise component of the velocity field (***Figure 2c***) was averaged over the span-wise direction, and also over the time period of 150 min.

### Cell growth measurement in U-shape traps in the EMPS

Here the choice of the strain, the medium, the culture condition and the instruments was the same as those for the measurement with the PDMS-based device, unless otherwise stipulated. Here, the cultured bacterial suspension was diluted to OD = 0.04 before it was inoculated on the coverslip of the EMPS. As sketched in ***Figure 2A***, the substrate consisted of drain channels (100 *μ*m wide, 7 mm long, 13 *μ*m deep) and U-shape traps (30 *μ*m wide, 80 *μ*m long, 1.0 *μ*m deep), which were prepared by the methods described in ref. (***Hashimoto et al. (2016)***). When the bilayer membrane was attached to the substrate, care was taken not to cover the two ends of the drain channel to use, so that cells in the drain could escape from it. After the assembly of the device with the bacterial suspension, it was fixed on the microscope stage inside the incubation box maintained at 37 °C. To fill the device with medium, we injected fresh medium stored at 37 °C from the inlet (***Figure 1–Figure Supplement 1***), at the rate of 60 ml/hr for 5 min by a syringe pump (NE-1000, New Era Pump Systems).

During the observation, the flow rate of the M9(Glc+a.a. and BSA) medium was set to be 2 ml/hr (approximately 0.2 mm/sec above the membrane), except that it was increased to 60 ml/hr (approximately 6 mm/sec above the membrane) at a constant interval, in order to remove the cells expelled from the trap efficiently. The time interval of this flush was 60 min before the observed trap was filled with cells, and 30 min thereafter.

### Evaluation of the rhodamine exchange efficiency in the EMPS

We used a non-motile mutant strain W3110 ΔfliC Δflu ΔfimA. Before the time-lapse observation, we inoculated the strain from a glycerol stock into 2 ml M9(Glc+a.a.) medium in a test tube. After shaking it overnight at 37 °C, we transferred 20 *μ*l of the incubated suspension to 2 ml fresh M9(Glc+a.a.) medium and cultured it until the OD at 600 nm wavelength reached 0.1-0.5. The bacterial suspension was finally diluted to OD = 0.1 before it was inoculated on the coverslip.

Before starting the observations, we injected a PBS solution with 10 *μ*M rhodamine to adsorb fluorescent dye on the surface of the coverslip, then removed non-adsorbed rhodamine molecules by injecting a pure PBS buffer. We repeated this medium exchange several times. For the observation, we used a laser-scanning confocal microscope, Nikon Ti2, equipped with a 100x (N.A. 1.49) oil immersion objective and operated by Nikon NIS-Elements. The resolution along the Z-axis was 0.12 *μ*m, and cross-sectional images were taken over a height of 10 *μ*m, at the time interval of 1.8 sec. In the presence of bacterial cells, we first monitored the fluorescent intensity while we switched the medium to supply from PBS without rhodamine to that containing 10 *μ*M rhodamine, at the flow rate of 6 mm/sec (approximately 60 ml/hr above the membrane) (***Figure 3bd***). The medium switch was performed by exchanging the syringe connected to the device. After this observation, followed by a time interval of a few minutes without flow of the solution in the device, we started to monitor the fluorescent intensity while we switched the medium from the PBS with rhodamine to that without rhodamine, at the flow rate of 6 mm/sec (***Figure 3ce***).

### Observation of the bacterial reductive division

We used a wild-type strain MG1655. Before the time-lapse observation, we inoculated the strain from a glycerol stock into 2 ml growth medium in a test tube. The same medium as for the main observation was used (LB broth, M9(Glc+a.a.) or M9(Glc)). After shaking it overnight at 37 °C, we transferred 20 *μ*l of the incubated suspension to 2 ml fresh medium and cultured it until the OD at 600 nm wavelength reached 0.1-0.5. The bacterial suspension was finally diluted to OD = 0.05 before it was inoculated on the coverslip.

For this experiment, we used a substrate with wells of 55 *μ*m diameter and 0.8 *μ*m depth. The well diameter was chosen so that all cells in the well can be recorded. The device was placed on the microscope stage, in the incubation box maintained at 37 °C. The microscope we used was Leica DMi8, equipped with a 100x (N.A. 1.30) oil immersion objective and operated by Leica LasX. To fill the device with growth medium, we injected fresh medium stored at 37 °C from the inlet (***Figure 1–Figure Supplement 1***), at the rate of 60 ml/hr for 5 min by a syringe pump (NE-1000, New Era Pump Systems).

In the beginning of the observation, growth medium was constantly supplied at the rate of 2 ml/hr (flow speed approximately 0.2 mm/sec above the membrane). When a microcolony composed of approximately 100 cells appeared, we quickly switched the medium to a non-nutritious buffer (PBS or M9 medium with *α*-methyl-D-glucoside (*α*MG), see ***Table 1***) stored at 37 °C, by exchanging the syringe. The flow rate was set to be 60 ml/hr for the first 5 minutes, then returned to 2 ml/hr. Throughout the experiment, the device and the media were always in the microscope incubation box, maintained at 37 °C. Cells were observed by phase contrast microscopy and recorded at the time interval of 5 min.

The cell volumes were evaluated as follows. We determined the major axis and the minor axis of each cell, manually, by using a painting software. By measuring the axis lengths, we obtained the set of the lengths *L* and the widths *w* for all cells. We then estimated the volume *v* of each cell, by assuming the spherocylindrical shape of the cell: 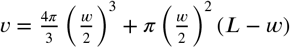. We estimated the uncertainty in manual segmentation at ±0.15 *μ*m.

### Simulations

Details on the derivation of the functional forms of *λ*(*t*) and *μ*(*t*) are provided in ***Appendix 1***. The parameters were evaluated as follows. First, from the observations of the exponential growth phase, we determined the growth rate *λ*_0_ directly. This allowed us to set the replication speed *μ*_0_ too, by using the relation 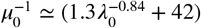 proposed by Wallden *et al*. (***Wallden et al. (2016)***) (the units of *λ*_0_ and *μ*_0_ are min^-1^). Concerning the volume threshold for initiating the replication, we found such a value of 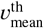 that reproduced the experimentally observed mean cell volume in the growth phase. The standard deviation 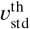. was set to be 10% of the mean 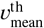, based on the relation found by Wallden *et al*. (***Wallden et al. (2016)***). They also measured the fluctuations of the time length of the C+D period; this led us to estimate 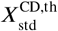 at 5% of 〈*X*^CD,th^〉, i.e., 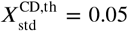. On the septum positions, we measured their fluctuations and found little difference in 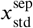 among the different growth conditions we used, and also in the non-nutritious case (***Figure 4–Figure Supplement 4***). We therefore used a single value 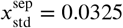 for all simulations.

Next we evaluated the parameters characterizing the time-dependent rates. The growth rate *λ*(*t*) can be determined independently of the cell divisions, because the total volume 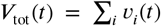 grows as 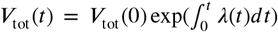. With *λ*(*t*) given by ***Equation 4***, we compared *V*_tot_(*t*) with experimental data and determined the values of *A* and *C* (***Figure 5c***). Finally, only *k* and *τ* in ***Equation 5*** remained as free parameters. We tuned them so that the mean cell volume *V*(*t*) and the number of the cells *n*(*t*) observed in the simulations reproduced those from the experiments (***Figure 5d***).

The parameter values determined thereby are summarized in ***Table 3***, for the simulations for LS→PBS and M9(Glc+a.a.)→PBS. Note that we did not perform simulations for M9(Glc)→M9(*α*MG), because we found it difficult to fit the experimental data for *V*_tot_(*t*) and *n*(*t*) in this case (***Figure 4–Figure Supplement 2***). This suggests that the consequence of the replacement of glucose by glucose analog *α*MG may not be simple starvation.

Concerning the initial conditions, we started the simulations from 10 cells with volumes in the range of 0.07-1.13 *μ*m^3^, randomly generated from the uniform distribution. The cells grew in the exponential phase (with the constant growth rate *λ*_0_ and the replication speed *μ*_0_) until the number of cells reached 100,000. We then randomly picked up 10 cells from this “precultured” sample and used them as the initial population of each simulation. Thus, the cell cycles of the cells were sufficiently mixed.

## Acknowledgments

We acknowledge discussions with H. Chaté, Y. Furuta, H. Nakaoka, T. Hiraiwa, Y. Kitahara and D. Nishiguchi. We thank I. Naguro for letting us use the OSMOMAT 030. This work is supported by KAKENHI from Japan Society for the Promotion of Science (JSPS) (No. 16H04033, No. 19H05800), a Grant-in-Aid for JSPS Fellows (No. 20J10682) and by the grants associated with the “Planting Seeds for Research” program and Suematsu Award from Tokyo Tech.

## Appendix 1 Derivation of the functional forms of *λ*(*t*) and *μ*(*t*)

Here we describe how we obtained the functional forms of *λ*(*t*) and *μ*(*t*), ***Equation 4*** and ***Equation 5***, which were used to define our sCH model for the bacterial reductive division. We consider the situation where growth medium is switched to non-nutritious buffer at *t* = 0; therefore, *t* denotes time passed since the switch to the non-nutritious condition. First, on the basis of the Monod equation (***Monod (1949)***), we assume that the growth rate *λ*(*t*) can be expressed as a function of the concentration of substrates, *S* (*t*), inside each cell:

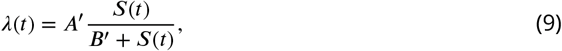

where *A′* and *B′* are constant coefficients. We consider that substrates in each cell are simply diluted by volume growth and consumed at a constant rate *C*. Therefore,

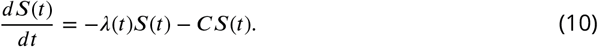

Note that there is no uptake of substrates because of the non-nutritious condition considered here. By combining ***Equation 9*** and ***Equation 10***, we obtain the following differential equation for *S*(*t*):

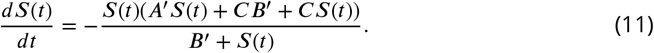

We can solve this equation and obtain

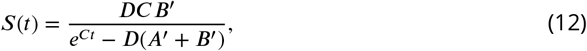

where *D* is a constant of integration. Substituting it to ***Equation 9***, we obtain

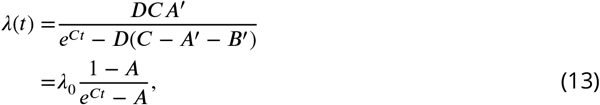

with *A* = *D*(*C* – *A′* – *B′*) and *λ*_0_ = *λ*(0) = *DCA*^1^ /(1 – *D*(*C* – *A′* – *B′*)) being the growth rate in the growth phase, before the onset of starvation.

Next, we determine the functional form of the replication speed *μ*(*t*), more precisely the progression speed of the C+D period. We first note that the C+D period mainly consists of DNA replication, followed by its segregation and the septum formation (***Wang and Levin (2009)***). Most parts of those processes involve biochemical reactions of substrates, such as deoxynucleotide triphosphates for the DNA synthesis, and assembly of macromolecules such as FtsZ proteins for the septum formation. We therefore consider that the replication speed is determined by the intracellular concentration of relevant substrates and macromolecules. Here we simply assume that the progression of the C+D period can be represented by assembly-like processes of relevant molecules, represented collectively by RM_C+D_. We then consider that the progression speed *μ*(*t*) is given through the Hill equation, which usually describes the binding probability of a receptor and a ligand, with cooperative effect taken into account. Specifically,

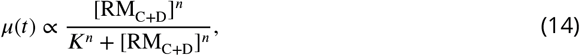

where *n* is the Hill coefficient, and *K* is the equilibrium constant of the (collective) assembly process. One can evaluate the time evolution of the concentration [RM_C+D_] in the same way as for *S*(*t*). With a constant consumption rate *E*, the differential equation can be written as

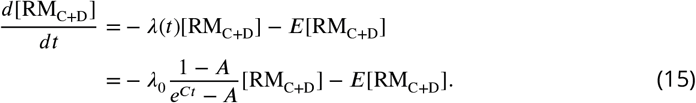

The solution to this equation is

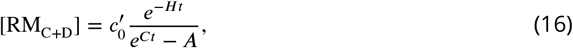

with a constant 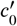 and *H* = *E* – *λ*_0_(1 – *A*)/*A*. To reduce the number of the parameters in the model, we simply assume that [RM_C+D_] exponentially decreases during starvation, *i.e*.,

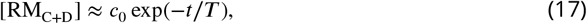

with initial concentration *c*_0_ and the degradation time scale *τ*. As a result, we obtain

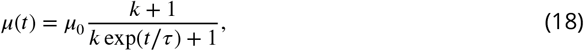

with *k*: = (*K/c*_0_)^*n*^, *τ* = *T/n*, and *μ*_0_(= *μ*(0)) being the replication speed in the growth phase before the onset of starvation.

## Appendix 2 Theoretical conditions for the scale invariance

Here, we describe the detailed derivation of the sufficient condition for the scale invariance of the cell size distribution. We start from the time evolution equation, ***Equation 6***:

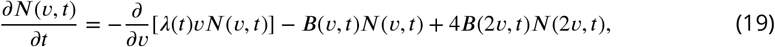

where *λ*(*t*) is the growth rate and *B*(*v,t*) is the division rate function (see the main article for its definition). We assume the scale invariance in the form of ***Equation 2*** for the cell size distribution *p*(*v,t*) = *N*(*v,t*)/*n*(*t*), where 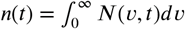 is the total number of the cells. For *N*(*v,t*), this reads

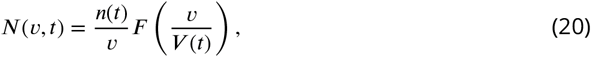

where *V*(*t*) is the mean cell volume at time *t, V*(*t*) = 〈*v*〉. One can then rewrite the time evolution equation (***Equation 19***) in terms of the function *F*(*x*) with *x* = *v*/*V*(*t*), as follows:

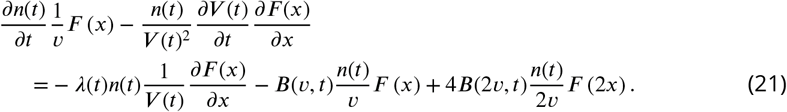

If we neglect cell-to-cell fluctuations of the growth rate, one can evaluate *λ*(*t*) from the total biomass growth, as follows:

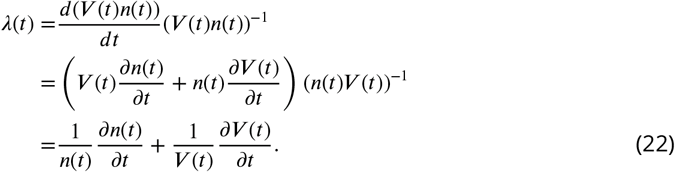

From ***Equation 21*** and ***Equation 22***, we obtain

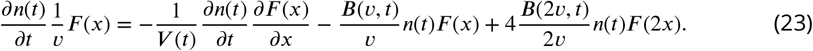

Now, the time derivative of *n*(*t*) can be calculated as

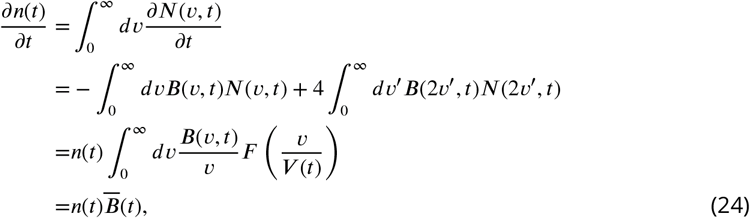

where 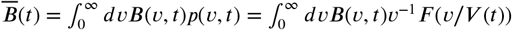. Substituting it to ***Equation 23***, we finally obtain the following self-consistent equation for *F*(*x*) (***Equation 7***):

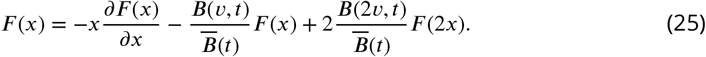

For the scale invariance, ***Equation 25*** should hold at any time *t*. In other words, the coefficient 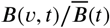 should be independent of both *t* and *V*(*t*). Note here that 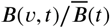 can be rewritten as

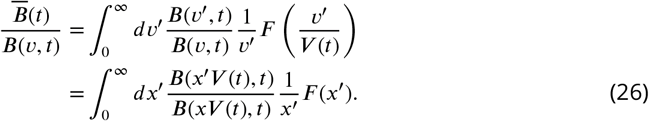

Therefore, 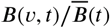 is time-independent if 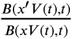 does not depend on *V*(*t*), being a function of two dimensionless variables *x* and *x′* only, as follows:

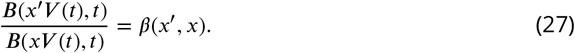

This condition can be rewritten as follows. For a constant *x*_0_, we can define *B_t_*(*t*) by

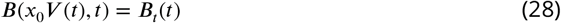

and *B_v_*(*x*) by

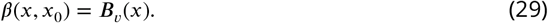

Then, the division rate can be expressed as *B*(*xV*(*t*), *t*) = *B_v_*(*x*)*B_t_*(*t*) for any *x* and *t*. This gives the sufficient condition we presented in the main text, ***Equation 8***,

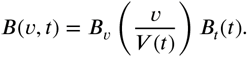

### Test of the derived conditions for *B*(*v, t*) and *F* (*x*)

We tested the sufficient condition for *B*(*v,t*) (***Equation 8***) and the resulting self-consistent equation for *F*(*x*) (***Equation 25***) with numerical data we obtained from our sCH model (***Figure 6d***). We evaluated the division rate *B*(*v,t*) in the simulations for LB → PBS, by measuring the number of division events of cells between time *t* and *t* + Δ*t*, divided by the number of cells at time *t*, where only the cells of volume between *v* and *v*+Δ*v* were considered in both the numerator and the denominator. Here, Δ*v* was set to be approximately 0.2 × *V*(*t*) and Δ*t* to be approximately 20-30 min for each time point, respectively. The value of *B*(*v*, 0) was determined by counting all division events in the exponential growth phase (*t* < 0). The ratio *B*,(0)/*B*,(*t*) can be evaluated by 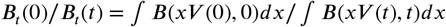. We found that the curves *B*(*v, t*)*B*,(0)/*B*,(*t*) taken at different *t* overlap reasonably well (***Figure 6d***, Inset), which support the variable separability condition of the division rate, ***Equation 8***.

Given the functional form of *B*(*v,t*) that we obtained numerically, we can also test the self-consistent equation for *F*(*x*), ***Equation 25***. Here we remind that the ratio 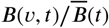 in ***Equation 25*** can be expressed as ***Equation 26***, and 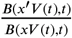 is time-independent (cf. ***Equation 27***). Therefore,

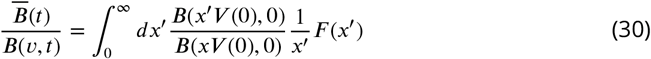

with *B*(*xV*(0), 0) = *B*(*v*, 0) = *B*(*v,t*)*B*,(0)/*B*,(*t*). Since we already confirmed the time independence of *B*(*v,t*)*B*,(0)/*B*,(*t*) (***Figure 6d***, Inset), we took the average of this quantity obtained at *t* = 0,30,60,90 min. Since the observed range of *v* is finite and *F*(*x*) almost vanishes for *x* > 2, we evaluated the integral over *x′* in the range 0 ≤ *x′* ≤ 2.

With the 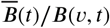 evaluated thereby, we substituted *F*(*x*) obtained by the simulations for LB→PBS to ***Equation 25*** (***Figure 6–Figure Supplement 2a***), using the time average of *F*(*x*) (the dashed line in ***Figure 6c***) and *F*(*x*) = 0 for *x* ≥ 2. We also tested ***Equation 25*** with *F*(*x*) obtained in the experiment for LB→PBS, in the same way as for the model (***Figure 6–Figure Supplement 2b***). In both cases, the right-hand side (rhs) of ***Equation 25*** differs significantly from the observed form of *F*(*x*).

### Theory with septum fluctuations

As a possible improvement of our theory, here we take into account septum fluctuations. We define the kernel function *q*(*ν*|*v*) which represents the probability that a mother cell of volume *v* produces daughter cells of volume *v* and *ν* – *v* (therefore, *q*(*ν*|*v*) = *q*(*v* – *ν*|*v*)). The time evolution equation of *N*(*ν,t*) is then written as

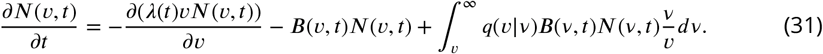

Now, assuming that the fluctuations of the septum position are Gaussian for simplicity, we have

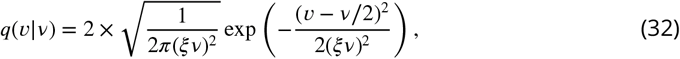

where the coefficient 2 corresponds to the two daughter cells produced from a single division event. Here, following experimental observations (***Figure 4–Figure Supplement 4***, see also (***Taheri-Araghi et al. (2015)***)), we assumed that the standard deviation is proportional to the volume of the mother cell, *v*, with coefficient *ξ* = 0.0325.

Under this modification, we obtain the self-consistent equation for *F*(*x*) as follows. One can again calculate the time derivative of *n*(*t*) as

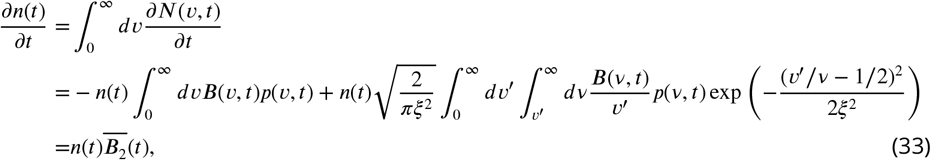

where 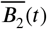 is defined by

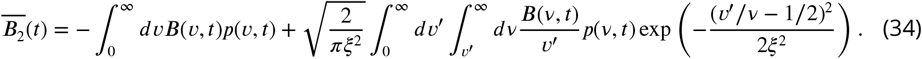

With *x* = *ν*/*V*(*t*) and *y* = *v*/*V*(*t*), one can obtain the self-consistent equation for *F*(*x*) as

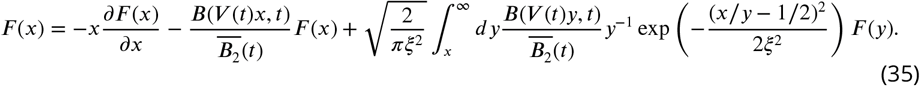

with

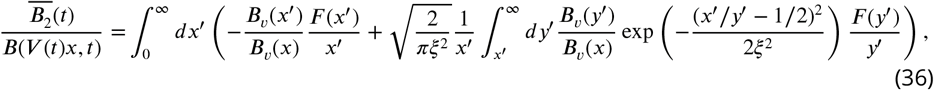

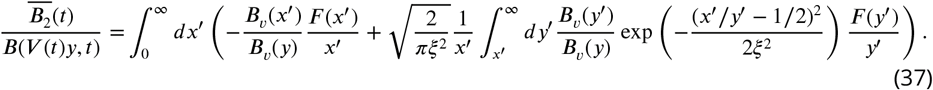

We tested this for both experimentally and numerically observed *F*(*x*), but the modification did not improve the results (***Figure 6–Figure Supplement 2***).

---

**Figure 1–Figure supplement 1.**
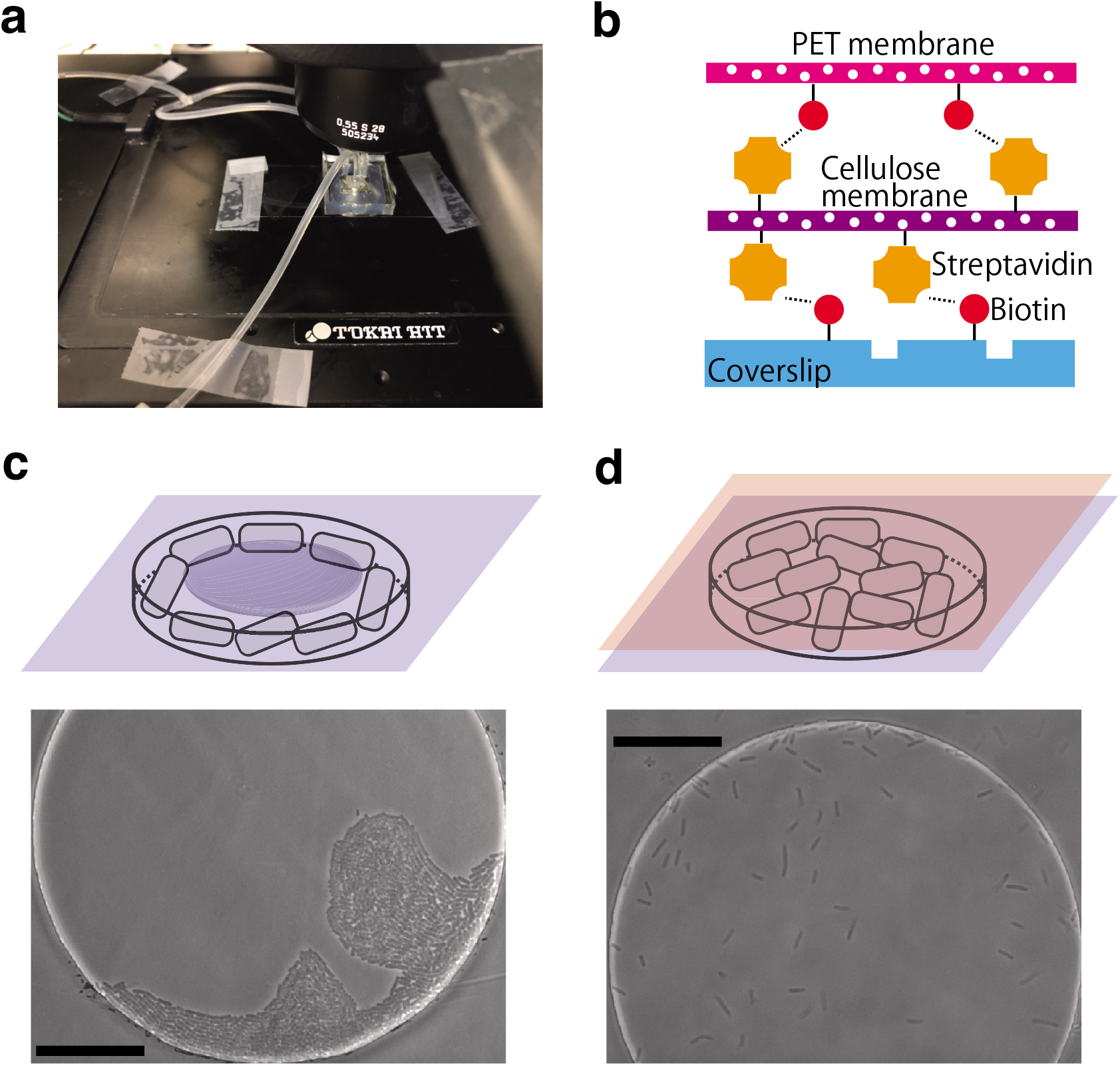
Supplementary figures on the setup of the EMPS. **a** Photograph of the device mounted on the microscope stage. The coverslip and tubes are fixed to the stage by mending tapes. **b** Illustration of the chemical bonding between a bilayer membrane and a glass coverslip. A streptavidin-decorated cellulose membrane is sandwiched by a biotin-coated PET membrane and coverslip. **c**(top) Sketch of growth of bacterial cells inside a circular well covered only by a cellulose membrane. Since the membrane bends and presses cell beneath, cells do not swim but form clusters, extending toward the wall. (bottom) Photograph of motile *E. coli* RP437 in a well (diameter 110 *μ*m, depth 1.1 *μ*m) covered only by a cellulose membrane. TB medium was constantly supplied at 37 °C (see Materials and Methods for details). Despite the motility, cells were confined near the wall and unable to swim freely (see also **Video 1**). The scale bar is 30 *μ*m. **d**(top) Sketch of growth of cells inside a well covered by a PET-cellulose bilayer membrane. The rigid bilayer membrane is sustained without bending, leaving a sufficient gap beneath for cells. (bottom) Photograph of motile *E. coli* RP437 in a well (same diameter and depth as in (**c**, bottom) covered by a bilayer membrane. Cells were able to swim freely (see also **Video 2**). Growth conditions are same as in (**c**, bottom).

**Figure 2–Figure supplement 1.**
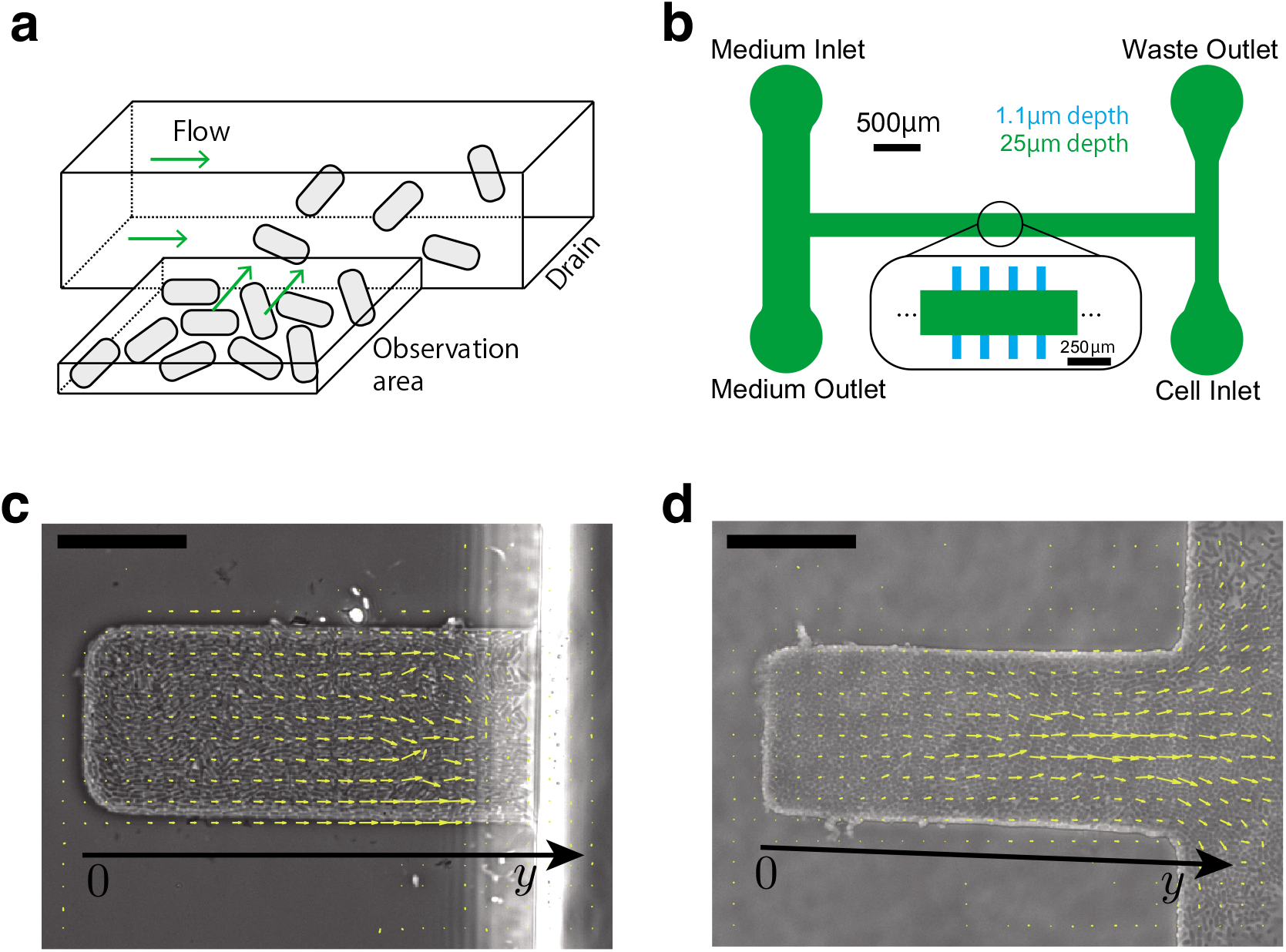
Supplementary figures on the setup of the EMPS. **a** Sketch of the design of microchannels in the PDMS-based device. Medium flow in the drain channel removes cells expelled from the observation area (trap). Nutrient is supplied to the cells inside the trap via diffusion from the drain channel. **b** The design of the PDMS-based device. The green region corresponds to the drain channel, and the blue regions are the U-shape traps. **c** Top view of the trap (30 *μm* wide, 88 *μm* long, 1.1 *μm* deep) in the PDMS-based device. The trap is filled with *E. coli* W3110 ΔfliC Δflu ΔfimA. The yellow arrows represent the velocity field of flow driven by cell proliferation, measured by particle image velocimetry (PIV). The scale bar is 25 *μ*m. See also **Video 3**. **d** Top view of the trap (30 *μ*m wide, 80 *μ*m long, 1.0 *μ*m deep) in the EMPS. See also **Video 4**.

**Figure 4–Figure supplement 1.**
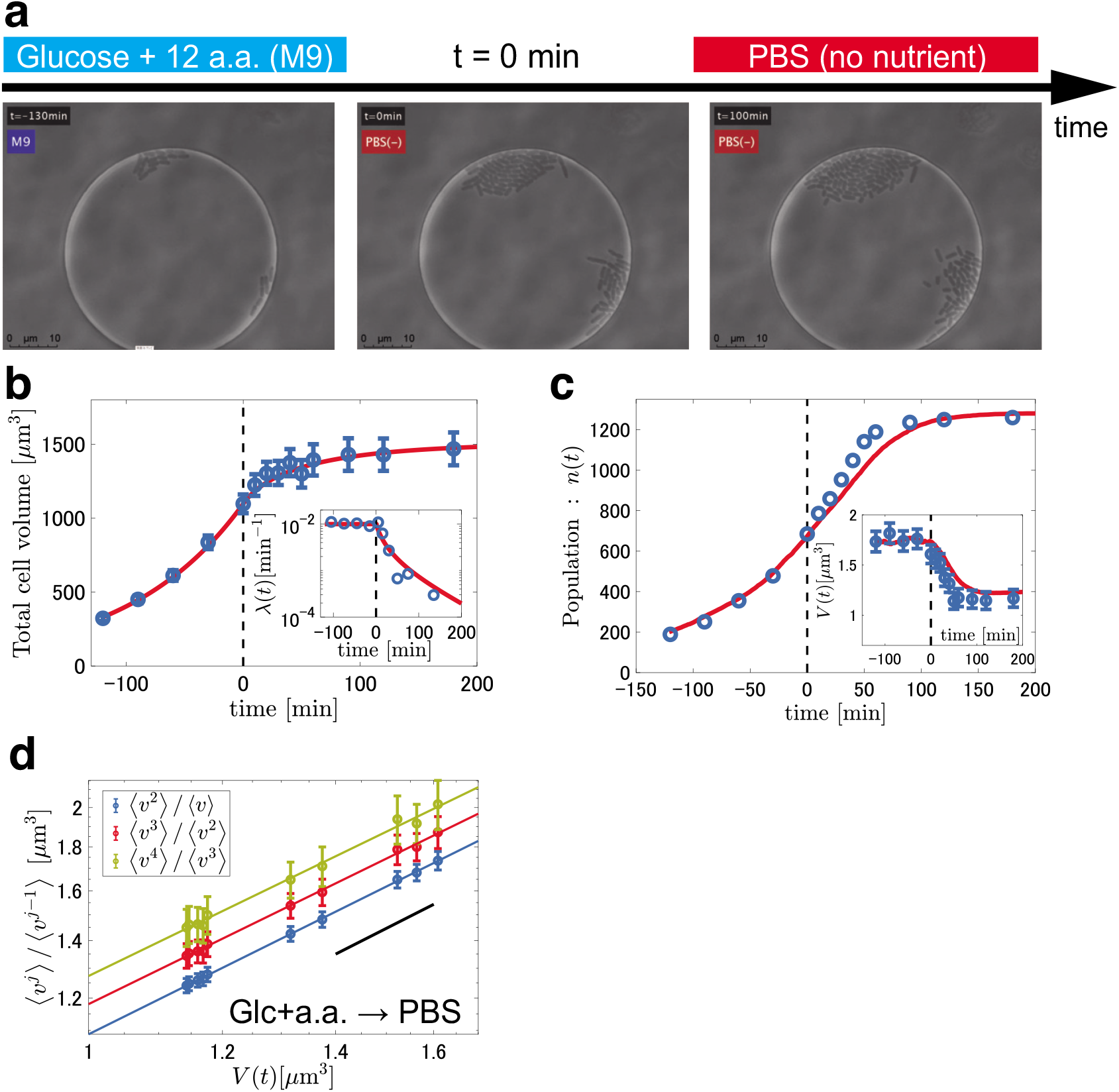
Results from the observations of reductive division in the case of M9(Glc+a.a.) → PBS. The data were collected from 17 wells. **a** Snapshots taken during the reductive division process. See also **Video 8**. **b,c** Experimental data (blue symbols) for the total cell volume *V_tot_*(*t*) **b**, the growth rate *λ*(*t*) (**b**, Inset), the number of the cells *n*(*t*) (**c**) and the mean cell volume *V* (*t*) (**c**, Inset) in the case of M9(Glc) → PBS, compared with the simulation results (red curves). The error bars indicate segmentation uncertainty in the image analysis (see Materials and Methods). *t* = 0 is the time at which PBS entered the device (black dashed line). **d** The moment ratio (*ν^j^*)/〈*v*^*j*−1^〉 against *V*(*t*) = 〈*v*〉. The error bars were estimated by the bootstrap method with 1000 realizations. The colored lines represent the results of linear regression in the log-log plots (see **Table 1** for the slope of each line). The black solid lines are guides for eyes indicating unit slope, i.e., proportional relation.

**Figure 4–Figure supplement 2.**
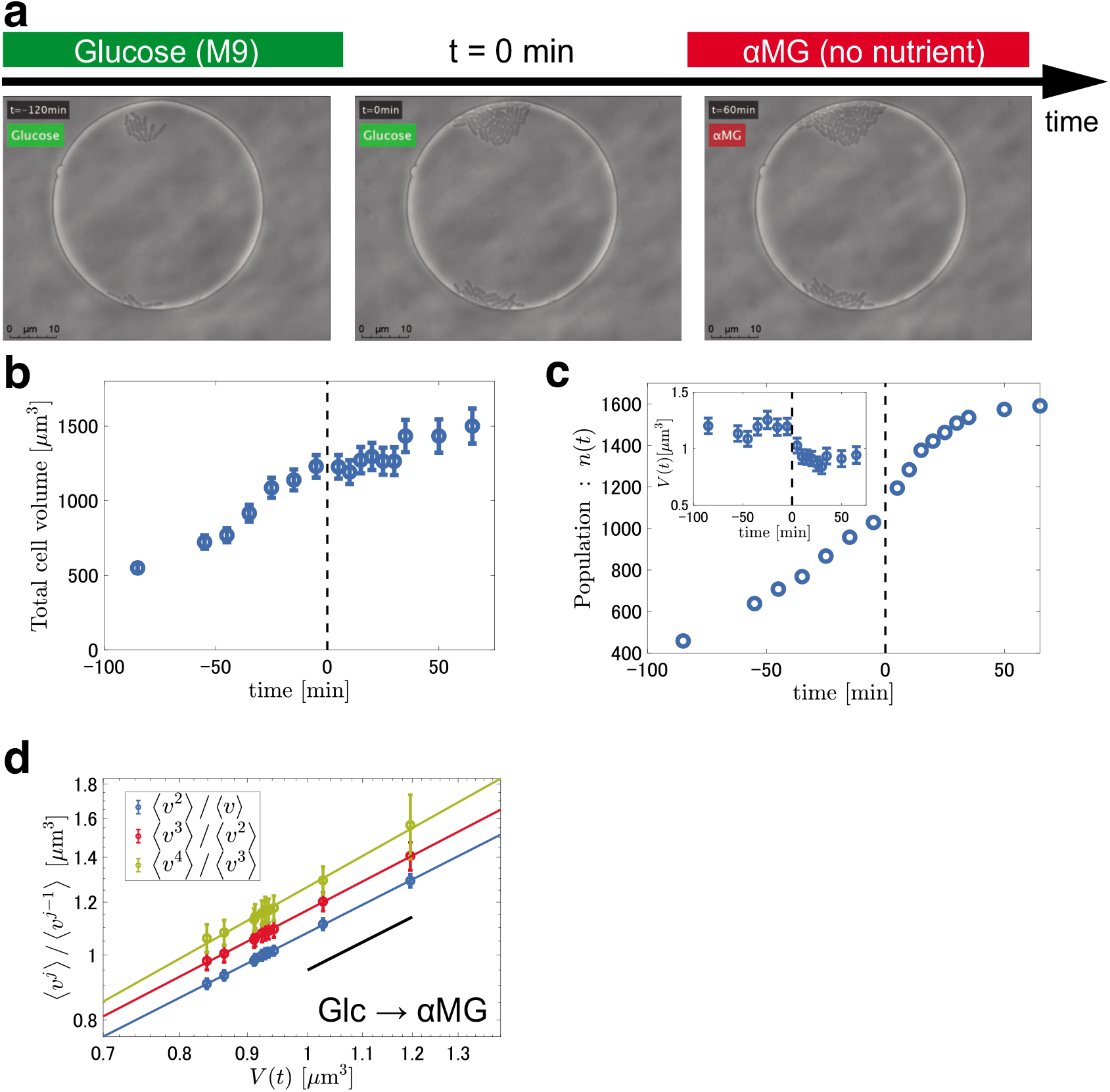
Results from the observations of reductive division in the case of M9(Glc) → M9(*α*MG). The data were collected from 26 wells. **a** Snapshots taken during the reductive division process. See also **Video 9**. **b,c** Experimental data for the total cell volume *V*_tot_ (*t*) (**b**), the number of the cells *n*(*t*) (**c**) and the mean cell volume *V*(*t*) (**c**, Inset) in the case of M9(Glc) → M9(*α*MG). The error bars indicate segmentation uncertainty in the image analysis (see Materials and Methods). *t* = 0 is the time at which *α*MG entered the device (black dashed line). **d** The moment ratio 〈*v^j^*〉/〈*v*^*j*−1^〉 against *V*(*t*) = (*v*). The error bars were estimated by the bootstrap method with 1000 realizations. The colored lines represent the results of linear regression in the log-log plots (see **Table 1** for the slope of each line). The black solid lines are guides for eyes indicating unit slope, i.e., proportional relation.

**Figure 4–Figure supplement 3.**
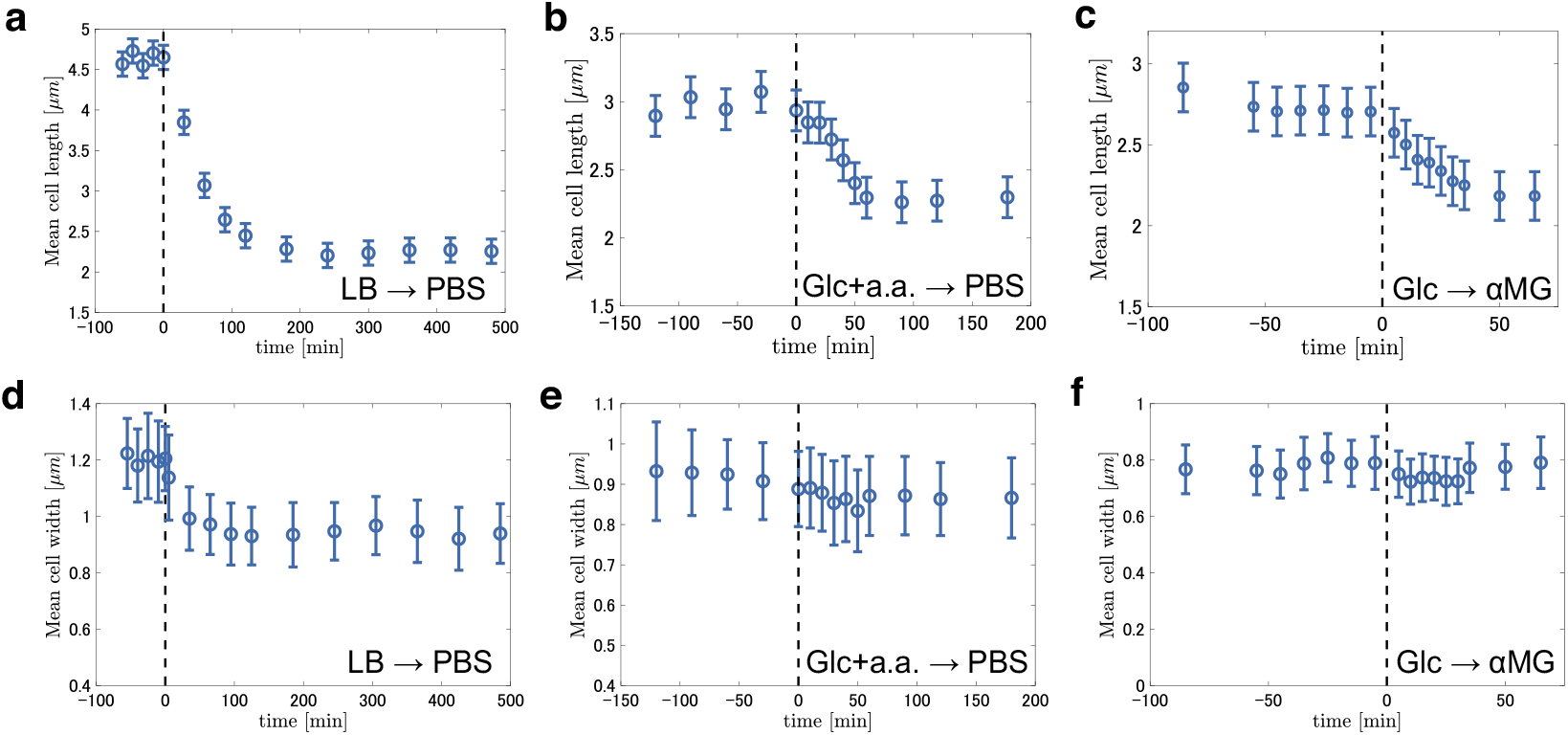
Time series of the mean cell length and the mean cell width during the starvation process. *t* = 0 is the time at which the non-nutritious buffer entered the device (black dashed line). **a**,**b**,**c** Time series of the mean cell length in the case of LB → PBS (**a**), M9(Glc+a.a.) → PBS (**b**) and M9(Glc) → M9(*α*MG) (**c**). The error bars indicate segmentation uncertainty in the image analysis (see Materials and Methods). **d,e,f** Time series of the mean cell width in the case of LB → PBS (**d**), M9(Glc+a.a.) → PBS (**e**) and M9(Glc) → M9(*α*MG) (**f**). The error bars indicate the standard deviation in each ensemble.

**Figure 4–Figure supplement 4.**
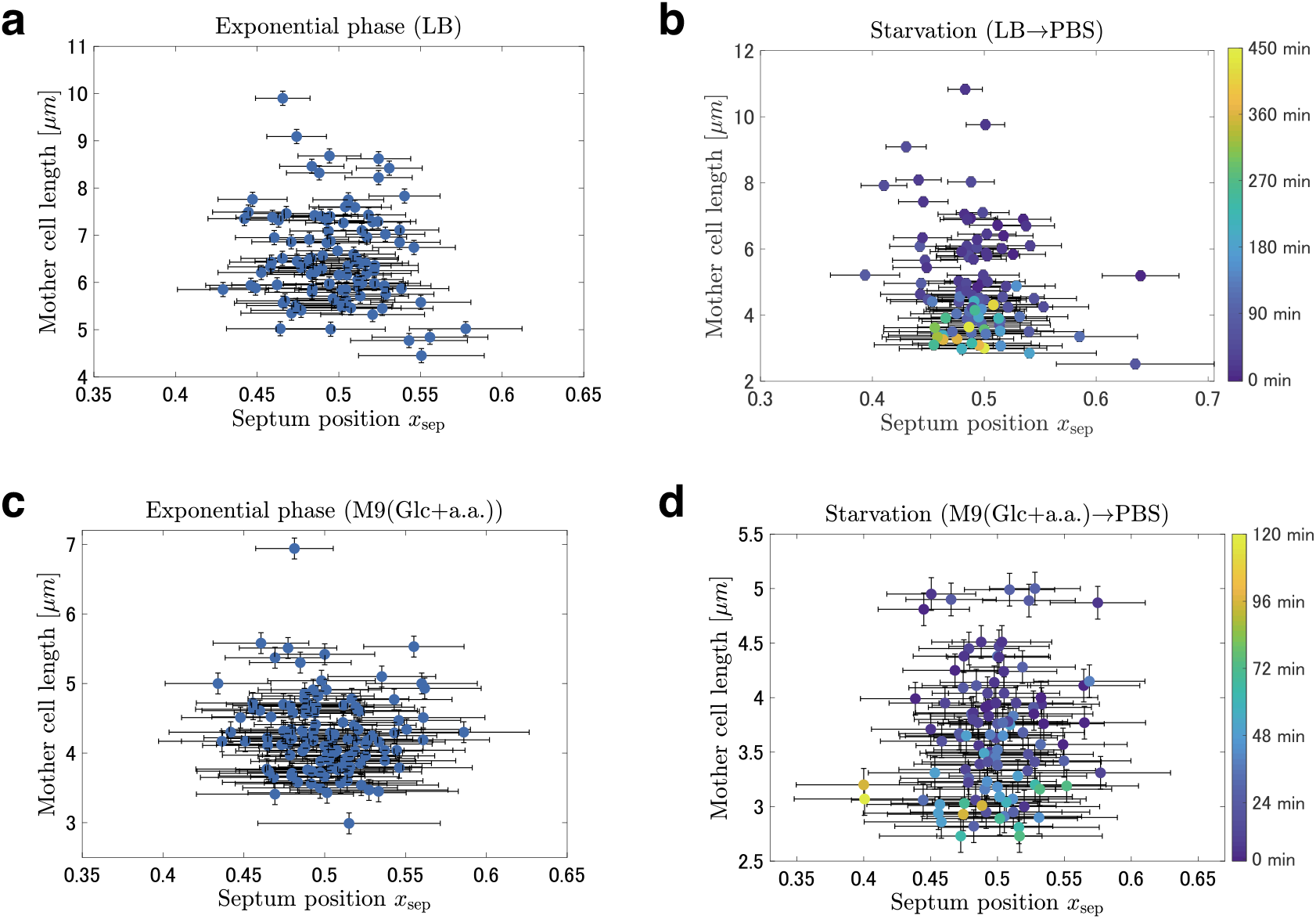
Cell-to-cell fluctuations of the septum position. Scatter plots of the mother cell length against the septum position *x*^sep^ are shown for four different cases. The data were taken from more than 100 cell division events chosen randomly for each case. The error bars indicate segmentation error in the image analysis. There is no visible correlation between the mother cell size and the standard deviation of the septum position. **a** Scatter plot for the exponential growth phase in LB broth. The standard deviation of the septum position is 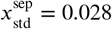. **b** Scatter plot during the starvation process for LB → PBS. The color represents the time passed since PBS entered the device. The standard deviation of the septum position is 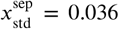. **c** Scatter plot for the exponential growth phase in M9(Glc+a.a) medium. The standard deviation of the septum position is 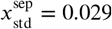. **d** Scatter plot during the starvation process for M9(Glc+a.a.) → PBS. The standard deviation of the septum position is 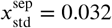.

**Figure 5–Figure supplement 1.**
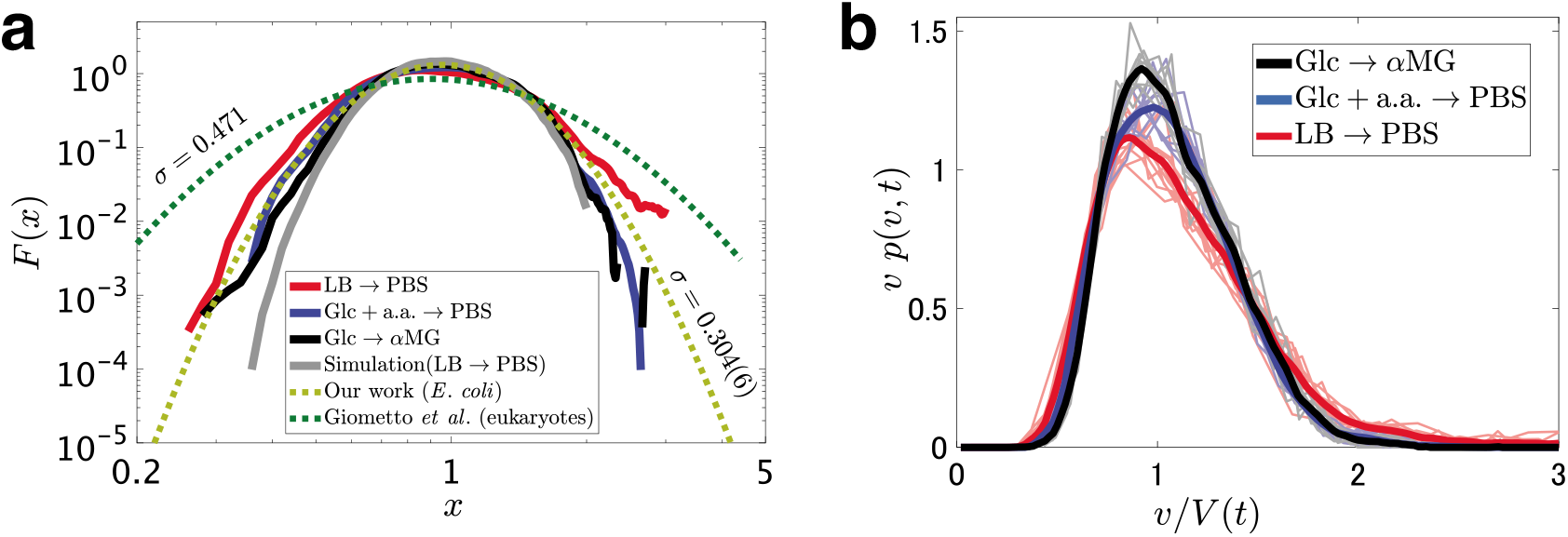
Comparison of the function *F* (*ν*/*V* (*t*)) = *νp*(*ν,t*) among different cases. **a** *F*(*x*) obtained by our experiments and simulation, as well as that obtained by Giometto *et al*. (***Giometto et al. (2013)***) for unicellular eukaryotes. The solid lines represent time-averaged data. The yellow dashed line is obtained by fitting **Equation 3** for the log-normal distribution to our experimental data. The green dashed line is the fitting result by Giometto *et al*. (***Giometto et al. (2013)***) for unicellular eukaryotes. *σ* is the standard deviation parameter of the log-normal distribution (see **Equation 3**). **b** *νp*(*ν,t*) for the three cases studied in this work, plotted in a linear scale. The raw data obtained at different times are shown by thin lines with relatively light colors, and the time-averaged data are shown by the bold lines. Instantaneous distributions (thin lines) also seem to be slightly but significantly different among the three cases.

**Figure 6–Figure supplement 1.**
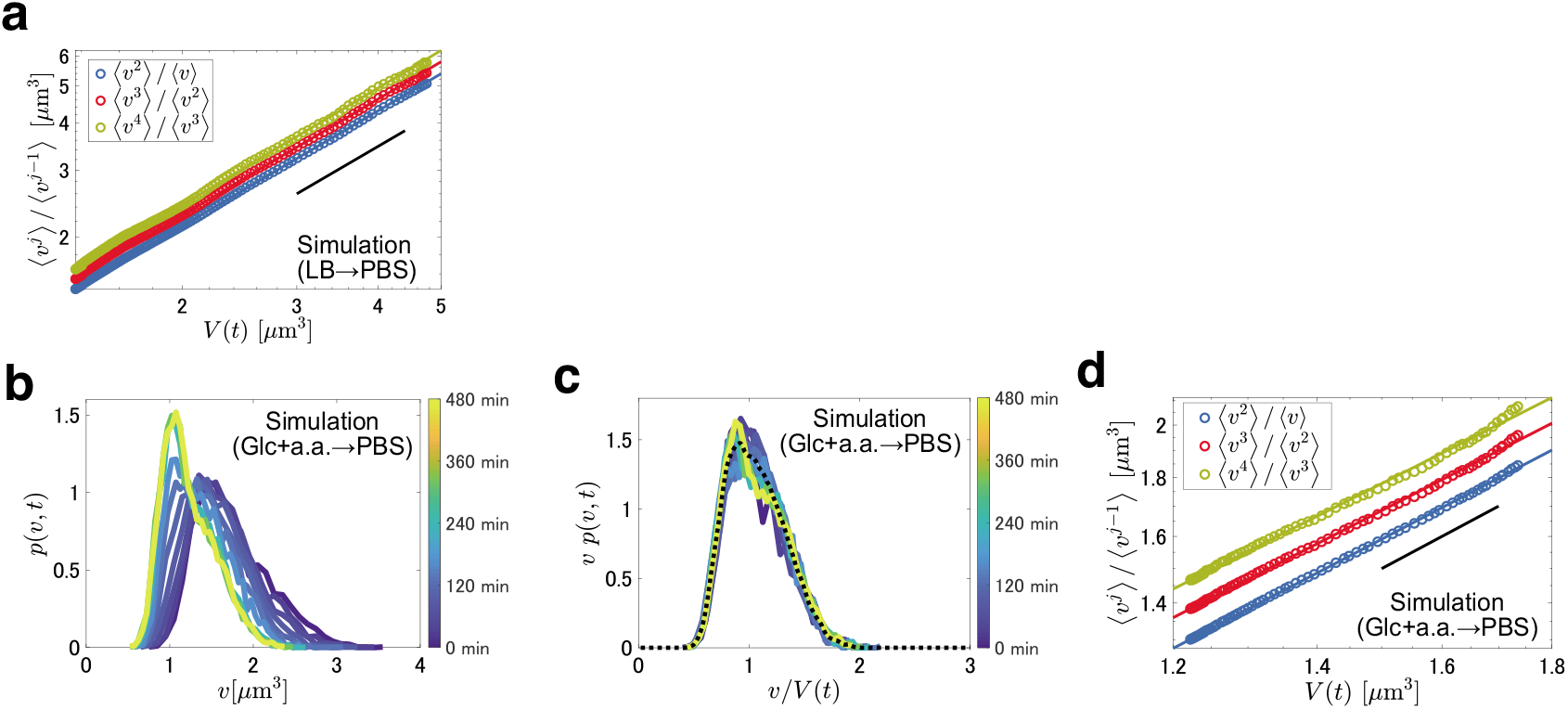
Supplementary figures on the simulation results. **a** The moment ratio 〈*v*〉/〈*ν*^*j*−1^)〉 against *V*(*t*) = 〈*ν*〉, in the model for LB → PBS. The error bars estimated by the bootstrap method with 1000 realizations were smaller than the symbol size. The colored lines represent the results of linear regression in the log-log plots (see **Table 1** for the slope of each line). The black solid lines are guides for eyes indicating unit slope, i.e., proportional relation. **b** Time evolution of the cell size distributions during starvation in the model for M9(Glc+a.a.) → PBS, obtained at *t* = 0,10,20,30,40,50,60,90,120,180 min from right to left. **c** Rescaling of the data in **b**. The overlapped curves indicate the function *F*(*ν*/*V*(*t*)) in **Equation 2** in the main text. The dashed line represents the time average of the datasets. **d** The moment ratio 〈*ν^j^*〉/〈*ν*^*j*−*i*^) against *V* (*t*) = 〈*v*〉, in the model for M9(Glc+a.a.) → PBS. The error bars estimated by the bootstrap method with 1000 realizations were smaller than the symbol size.

**Figure 6–Figure supplement 2.**
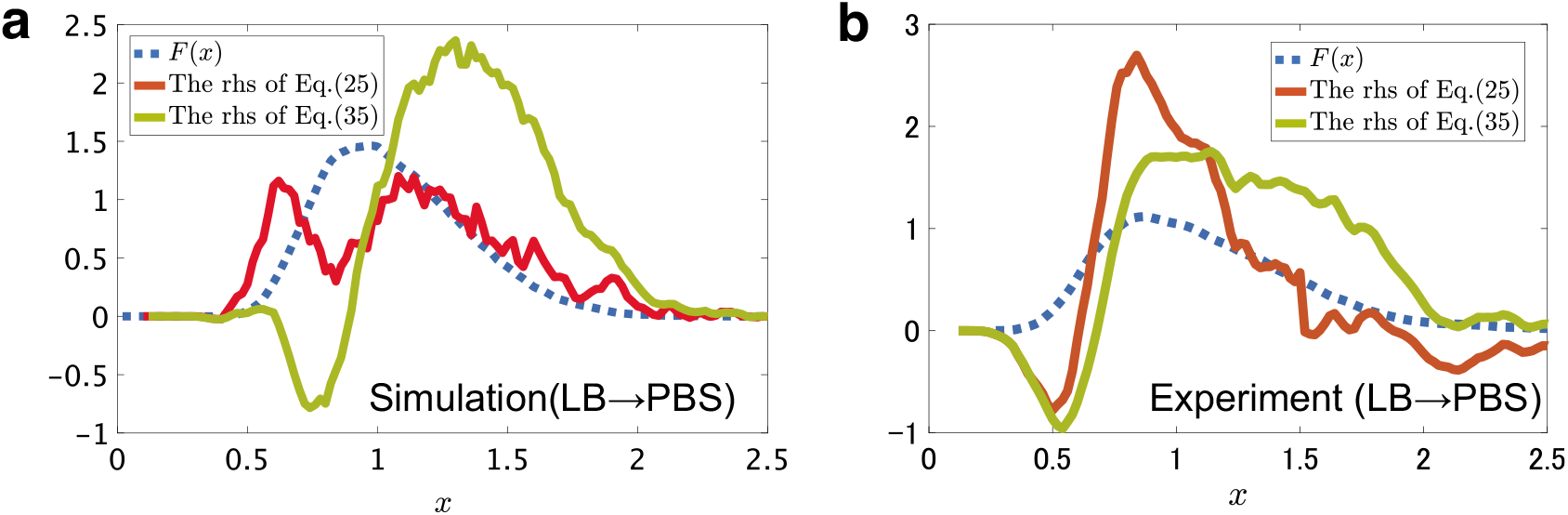
Test of the self-consistent equations derived in **Appendix 2**. The functional form of *F*(*x*), obtained numerically (**a**) or experimentally (**b**) for the case LB → PBS, is compared with the right-hand side (rhs) of the self-consistent equations derived in **Appendix 2**, **Equation 25** and **Equation 35**. The effect of septum fluctuations is neglected in **Equation 25** and considered in **Equation 35**. *F*(*x*) is obtained as the time average of the instantaneous data.

